# Phosphoproteomic analysis of the AKT signalling axis in cutaneous squamous carcinoma progression reveals novel therapeutic targets

**DOI:** 10.1101/2022.10.03.510591

**Authors:** R Button, C Harwood, RFL O’Shaughnessy

**Author notes:** Address for Correspondence: Ryan O’Shaughnessy, Centre for Cell Biology and Cutaneous Research, 4 Newark Street, London, E1 2AT,.

## Abstract

Cutaneous Squamous Cell Carcinoma (cSCC) represents about 20% of all non-melanoma skin cancers. Whilst generally low risk to patients, metastases are associated with a poor prognosis. cSCC incidence is increasing, owing to an ageing population, greater exposure to UV radiation, and more patients receiving immunosuppressive treatments associated with organ transplants. Therefore, there is interest in identifying new biomarkers that may be to track progression of the disease and to exploit as therapeutic vulnerabilities. We show dynamic changes in AKT expression in precursor lesions and in SCC tumour tissue, with initial loss of AKT activity followed by progressive and widespread increase in AKT activity in SCC.

Phosphoproteomic analysis and kinase substrate enrichment analysis on a panel of isogenic cSCC cell lines representing different stages of the disease from premalignancy to metastasis revealed several up-regulated kinases and AKT-targets. From this analysis we chose DNA dependent protein kinase (DNA-PK), a key kinase upstream of AKT phoshorlyation, and N-Myc downstream-regulated gene 2 (NDRG2) a downstream AKT phosphorylation target, to investigate in further detail. Both proteins were up-regulated and mis-expressed in a panel of SCC tissue from different patients. We therefore explored the potential of inhibiting DNA-PK and NDRG2 as cSCC treatments. Treatment with the iron chelator Dp44mT decreased levels of phosphorylated NDRG2 and led to significant losses to viability and reduced migration in our cSCC cell lines, while DNA-PK inhibition promoted the differentiation of premalignant and early-stage SCC cell lines. Our results suggest that NDRG2 and DNA-PK may be viable targets in cSCC treatment, with effectiveness at different stages of SCC progression.

## Introduction

Cutaneous squamous cell carcinoma (cSCC) is the second most common form of non-melanoma skin cancer (NMSC), representing ∼20% of these lesions (Lomas et al., 2012). Key risk factors include UV radiation exposure, old age, and immunosuppression. The bulk of cases occur on sun-exposed regions of the body (Kosutic et al., 2019). The majority of cSCC originate from precursor lesions known as actinic keratoses (AKs) and Bowen Disease (Criscione et al., 2009; Kao, 1986). Given the ageing population, increased UV exposure, and the increase in patients on immunosuppressive regimens following procedures like organ transplantation, cSCC incidence is increasing (Rembielak & Ajithkumar 2019; Conforti et al., 2019). In the USA, an estimated 700,000 cases are diagnosed annually, with metastases from these accountable for between 3900-8800 deaths (Kauvar et al 2015; Sarin et al., 2020). Therefore, cSCC represents a significant health and economic burden (Vallejo-Torres et al., 2014).

Whilst in the majority of cases surgical resection is the preferred treatment option, due to the locations of the tumours, there are instances where this strategy has limited usability (Kosutic et al., 2019). Notably, around 5% of cSCC progress to an invasive form, whereupon the prognosis of the patient becomes considerably worse (Brantsch et al., 2008; Lambert et al., 2013). Currently there are no approved targeted inhibitors for metastatic cSCC, highlighting the interest in the identification and development of novel targeted treatment options.

A key feature of cSCC is that they carry a high mutational burden (Inman et al., 2018). Therefore, genomic based approaches of identifying novel drug targets can be difficult A more readily translatable option is to identify kinases and downstream kinase targets whose activity are increased in cSCC. Kinase activity is elevated in numerous cancers and can be easily inhibited by small molecule inhibitors (Knapp 2018). During SCC progression down-regulation of the skin barrier-associated AKT1 activity followed by the increase in malignancy-associated AKT2 activity (O’Shaughnessy et al, 2007a and b). The mechanism of this change in AKT isoform phosphorylation is not known to date but is a key component of SCC formation and progression. A significant gap in our knowledge, with both prognostic and therapeutic implications, is how in skin, different AKT isoforms can be differentially regulated, and how they regulate different molecular targets.

As the progression of cSCC is a multi-stage disease, understanding at which points changes occur may be crucial in understanding drivers of the disease, as well as identifying potential therapeutic targets. Recent studies have addressed this regarding the genomic and methylation profiles, by comparing healthy epidermal keratinocytes, precursor AK lesions, and cSCCs (Inman et al., 2018; Thomson et al., 2021; Hervas-Marin *et al*., 2019; Rodriguez-Paredes et al., 2018). However, only limited work has conducted on SCCs at a phosphoproteomic level (Tiwari et al., 2017)

In this study, we used a mass spectrometry-based phosphoproteomic analysis on a series of cSCC cell lines from the same patient, each representing a different stage of the disease progression, to generate a list of predicted elevated kinases, and downstream phosphotargets. Focusing on AKT signalling we confirm increased phosphorylation of both AKT and the DNA damage-related kinase PRKDC. In cSCC. PRKDC inhibition pushed cSCC lines towards terminal differentiation and reduced AKT phosphorylation, suggesting that PRKDC is an important upstream kinase of AKT. Phosphorylation of the AKT target N-Myc Downstream Regulator 2 (NDRG2) was increased in cell lines and cSCC samples. NDRG2 inhibition by iron chelation reduced cSCC viablility. In summary, phosphoproteomic analysis has identified key components of an AKT signalling axis in cSCC that can potentially be exploited for therapeutic benefit.

## Results

### The majority of phosphorylation changes occurred in the early stages of cSCC progression

We performed mass spectrometry based phosphoproteomic analysis on an isogenic series of cSCC lines, comprising of pM1 (pre-malignant), Met1 (initial cSCC occurrence), Met2 (recurrence post-resection of Met1), and Met4 (lymph node metastasis of Met2), which were all derived from the same immunosuppressed patient (Proby et al., 2000). The Met1, Met2, and Met4 lines have decreased keratin 10 expression, as well as increased vimentin expression and decreased E-Cadherin expression, suggesting impaired terminal differentiation and possible epithelial-to-mesenchymal transition (Yang et al., 2004; Winter et al., 1983) (Supplementary Figure S1A and B).

We followed this with a comparison of altered phosphoproteins between pM1 and Met1, Met2, and Met4, which modeled the change from a precursor lesion (pM1) to cSCC (Met1, Met2) through to metastasis (Met4) (Supplementary Table ST4). The numbers of substrates with increased phosphorylation were roughly equal between the lines (1120, 899 and 1261 for Met1, Met2 and Met4, respectively), with 524 shared across all three lines (Supplementary Figure S1C), indicating that a large proportion of phosphorylation changes are maintained throughout cSCC progression. Gene Ontology (GO) analysis revealed that the substrates increased in phosphorylation were involved in cell migration, growth factor signaling, and changes to actin filaments. Substrates that decreased in phosphorylation were related to various aspects of the cytoskeleton, keratinocyte differentiation, and cell cycle (Supplementary Figure S1D).

We performed Kinase Substrate Enrichment Analysis (KSEA), which is a predictive algorithm that estimates the changes to kinase phosphorylation based on detectable changes to their known substrates in our samples (Casado et al., 2013). Since kinases are generally readily druggable, we reasoned this could represent the most efficient method of identifying novel therapy routes. We performed KSEA on pM1 vs. Met1, Met2, and Met4. This generated a list of 20 hits that were elevated in at least one of the three cSCC lines (Figure 1A; Supplementary Table ST2). 70% (14/20) of the kinases were increased in activity according to KSEA in the Met1 line, with 6 of these shared with Met2 (12/20) and Met4 (6/20 total, all common with Met2) (Figure 1B). Several of our predicted kinases have been previously reported to have increased activity in cSCC, including AXL (Papadakis et al., 2011), and AKT (O’Shaughnessy *et al*., 2007). Together, our data suggest that most kinase activity changes happen at the initial stages of tumour onset.

**Figure 1:**
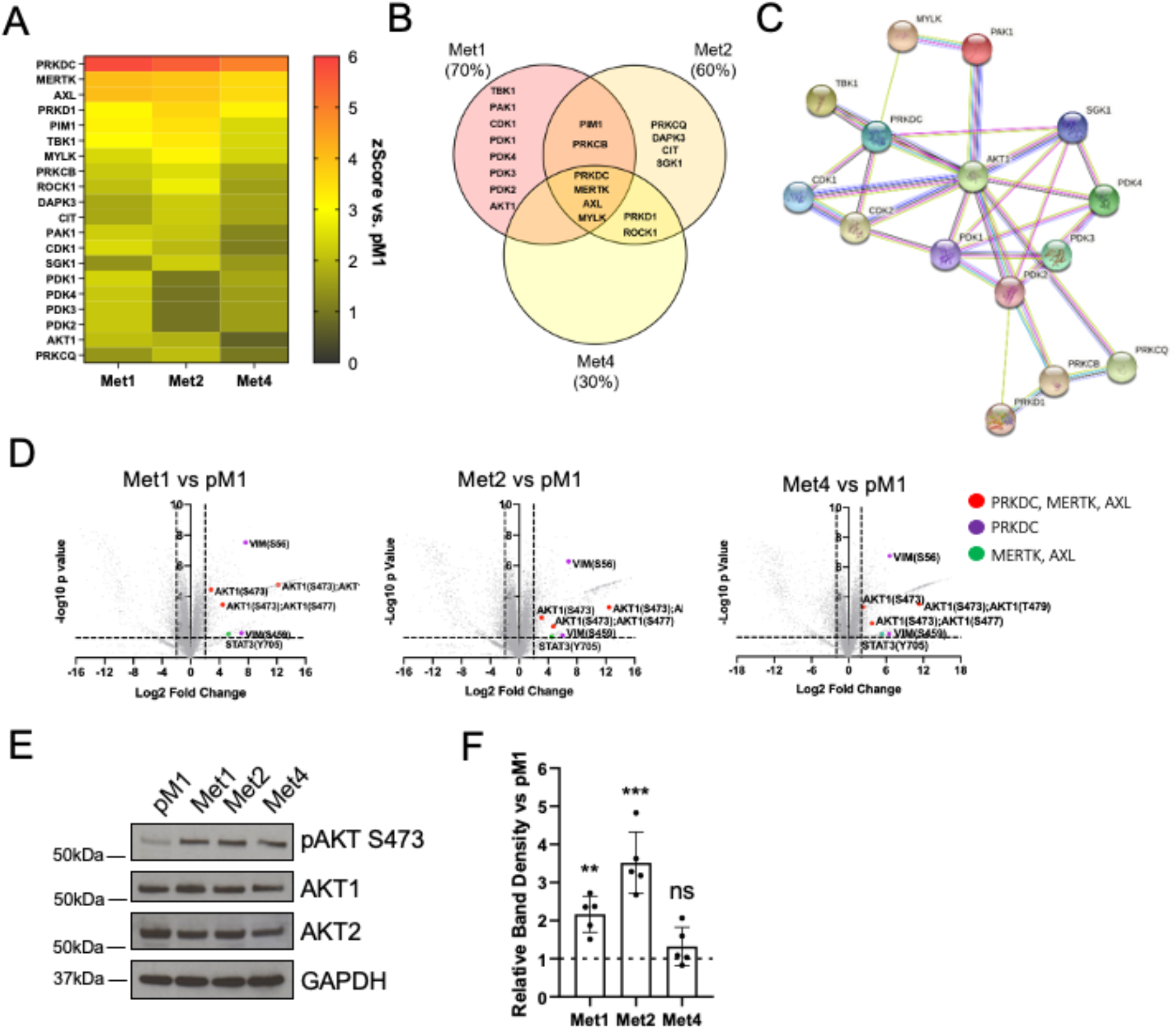
Phosphoproteomic analysis reveals PRKDC to be highly increased in phosphorylation in cSCC, with downstream links to AKT. **A**. Kinase Substrate Enrichment Analysis (KSEA) was applied to the dataset, with a heatmap displaying putative kinases that are significantly increased in at least one of the three lines (fold change ≥1.5, p≤0.05). **B**. Venn diagram showing the breakdown of the kinase list changes across the three cell lines. Percent values indicate proportion of total kinase list significantly increased in each line. **C**. STRING analysis of all kinase targets shows the putative protein-protein interactions that exist between them. **D**. Volcano plots showing all phosphoprotein substrates changed in (i) Met1, (ii) Met2, and (iii) Met4 vs. pM1. Dotted lines represent threshold limits, with 2/-2 for Log2 Fold Change, and 1.3 for –Log10 p value. Putative substrates for PRKDC, MERTK, and AXL based on the PhosphoSitePlus database are highlighted in the respective plots. **E**. Cell lines representing premalignant (pM1), initial cSCC (Met1) and recurrent cSCC (Met2) were treated with vehicle or NU7026 (NU) for 20hrs, Cells were harvested, and lysates analysed by immunoblot by probing as indicated, accompanying quantification of the mean relative band density of NU treatment vs the matching control is shown for pPRKDC and pAKT in each cell line (±SEM). GAPDH was used for loading control correction for pPRKDC, and total AKT for pAKT. Statistical analysis was performed by t-test for each cell line control vs treatment (** : P<0.01, *: P<0.05, ns: not significant, n=3 for each).

We formed a protein-protein interaction network of our list of kinases, revealing AKT to be a central kinase, interacting with 8 of the other candidates, and therefore potentially may represent a key signaling node in the changes throughout cSCC progression (Figure 1C). Consistent with this, we looked at the phosphorylated substrates of the three kinases with the largest activity increase based on KSEA - PRKDC, MERTK, and AXL - across our three cell lines. Most of the changes were associated with AKT modifications (Figure 1D). Notably, these were the only changes shared by all three kinases. AKT phosphorylation is altered in cSCC (O’Shaughnessy et al., 2007; Sully et al., 2013), with a switch from AKT1, which is important in epidermal barrier function (O’Shaughnessy et al., JBC, 2007a; Naeem et al.,2015; Naeem et al., 2017; Rogerson et al., 2021), to AKT2, which is associated with development and wound healing (Pankow *et al*., 2006). AKT phosphorylation was significantly increased in both the Met1 and Met2 lines (Figure 1E and F). As KSEA changes in Met4 were not significant, we focussed on the Met1 and Met2 lines for the remainder of the study.

To confirm our *in vitro* findings, we stained for pSerine 473 AKT (pAKT) in cSCC premalignancies (Actinic Keratoses (AKs) and Bowen’s Disease (BD)), cSCCs, and age-matched non-diseased skin, from both immunocompetent (IC) and immunosuppressed (IS) patients. Control skin showed discrete pAKT staining, largely limited to either the cornified layer of the epidermis (pAKT1 (O’Shaughnessy et al., 2007a and b)), with some also showing some basal activity (putative pAKT2 (O’Shaughnessy et al., 2007a and b)). Both AK and BD lesions showed more variability in localization and expression (Figure 2A). Almost all cSCC samples showed increased pAKT expression throughout the epidermis (Figure 2B). Stratifying AKs, BDs and cSCCs based on the immune status of the patient yielded no significant difference in pAKT staining, suggesting that increased AKT signalling was important in both immunocompetent and immunosuppressed tumour formation (Figure 2C).

**Figure 2:**
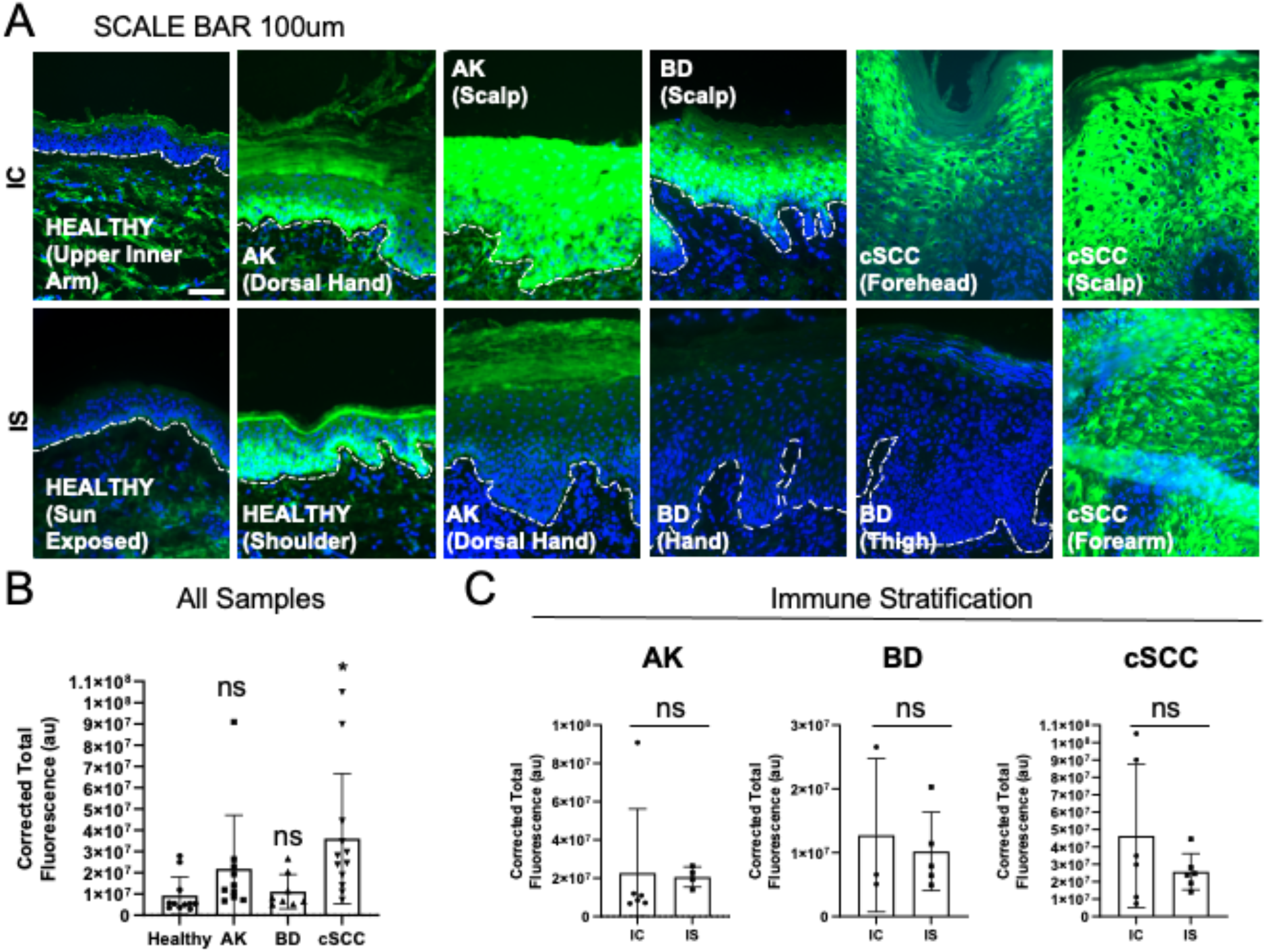
AKT phosphorylation is increased in cSCC and a subset of precursor lesions. **A**. Human tissue samples of Healthy Skin, Actinic Keratosis (AK), Bowen’s Disease (BD) and cSCC were stained with phospho-AKT antibody. Representative microscope images from both immunocompetent (IC) and immunosuppressed (IS) patients are displayed. Scale bar: 50um. **B**. The corrected total cell fluorescence of pAKT was calculated across the samples and presented as Healthy (n=11) vs AK (n=10), BD (n=8), and cSCC (n=12). Statistical Analysis was performed using ANOVA with Kruskal-Wallis as a post-test (*: P<0.05, ns: not significant). **C**. AK, BD and cSCC samples were stratified based on the patient immune status. Data are presented as mean ±sd (AK n=6 IC, n=4 IS; BD n=3 IC; n=5 IS; cSCC n=6 IC; n=6 IS), with statistical analysis performed by t-test.

### PRKDC phosphorylation increases with cSCC progression

Of our KSEA predicted kinases, PRKDC (also known as DNA-PK) showed the highest overall Z-score across all three cell lines. PRKDC is a serine/threonine kinase that has a role in the repair of double strand breaks to DNA following UV exposure, through the process of non-homologous end joining (NHEJ) (Yue et al., 2020). Aberrant expression of the kinase is associated with several tumours, particularly those with a high mutational load (Tan et al., 2020). Human epidermal keratinocytes exposed to varying UV doses show elevated PRKDC phosphorylation (Tu et al., 2013). PRKDC has also been linked to cSCC migration (Zhang et al., 2019), but little is known about its role in cSCC progression. AKT was directly linked to PRKDC in our protein-protein interaction network (Figure 1C) suggesting that it could contribute to the AKT changes we see in cSCC, and by extension, PRKDC inhibition could aid in restoring normal levels of AKT in the disease (Tu et al., 2013). Since AKT1 is important for normal epidermal homeostasis and epidermal barrier function, Pan-AKT inhibition could have unforeseen deleterious effects. As AKT2 is increased in phosphorylation in cSCC, it is possible that PRKDC specifically upregulates that isoform only, meaning targeting it could be a more precise approach to reducing cSCC-associated AKT activity.

To explore the phosphorylation of PRKDC in cSCC further, we stained our panel of healthy, AK, BD and cSCC patient samples with pPRKDC-T2609. Expression in healthy tissues was confined to the cornified layer, while AKs and BDs again showed a spectrum of expression towards positivity throughout the epidermis. In cSCC, there was strong, significantly increased pPRKDC expression throughout the epidermis (Figure 3A and B) (Figure 3B). Like pAKT, stratifying each of the lesion groups based on their immune status (IC vs IS) revealed no significant difference in pPRKDC signal (Figure 3C). Therefore, PRKDC phosphorylation increased over the course of disease progression, with the shift from a granular layer localization to a more widespread pattern throughout the epidermis.

**Figure 3:**
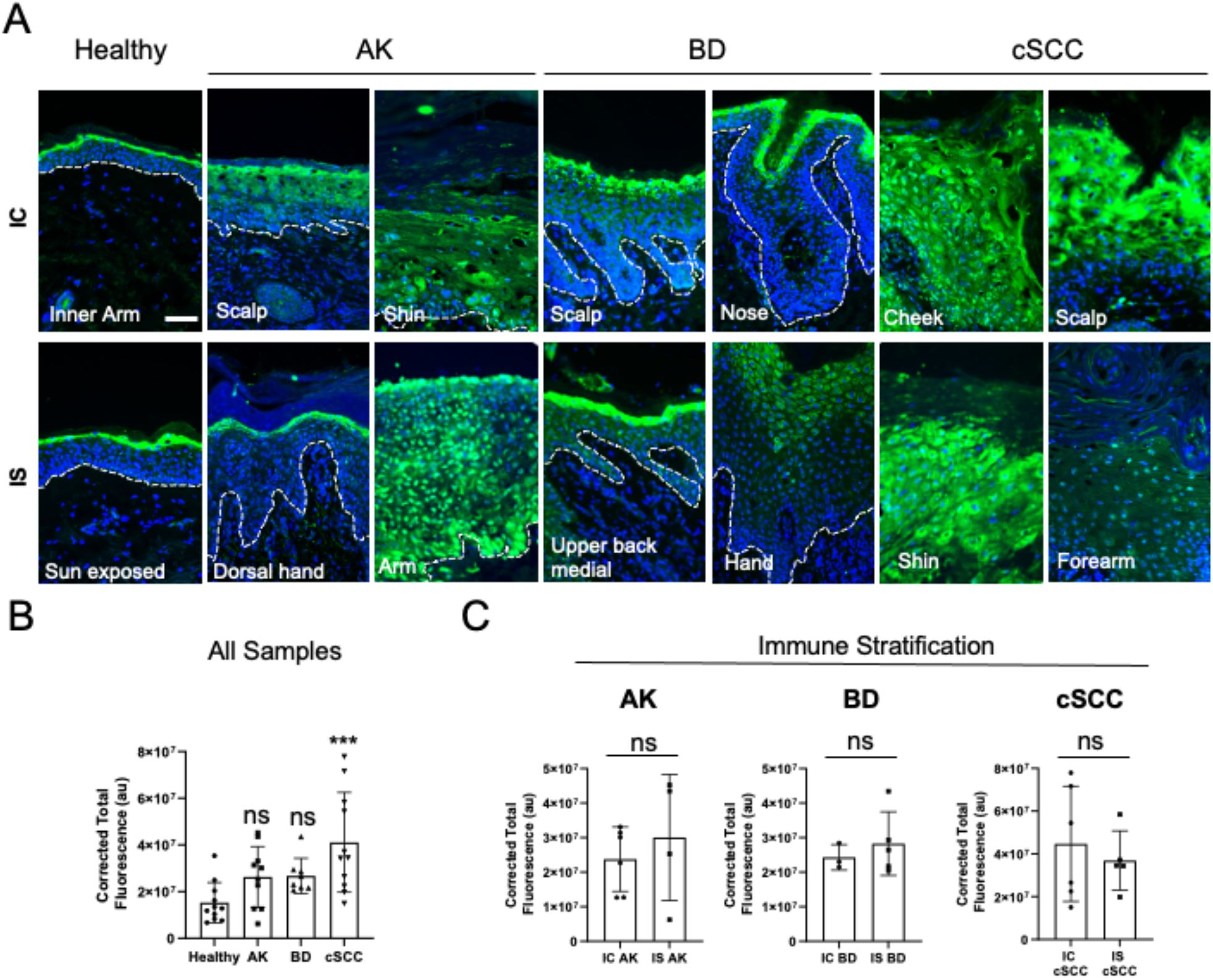
Expression of phosphorylated PRKDC increases with cSCC progression. **A**. Frozen human tissues representing Healthy, Actinic Keratosis (AK), Bowen’s Disease (BD) and cutaneous squamous cell carcinoma (cSCC) were stained for pPRKDC. Representative images taken with epifluorescence microscope are shown, with both immunocompetent (IC) and immunosuppressed (IS) tissues displayed. Scale bar: 50um. **B**. Corrected total fluorescence ±sd was calculated for each tissue (healthy n=11; AK n=10; BD n=8; cSCC n=11). Statistical Analysis was performed using ANOVA with Kruskal-Wallis as a post-test (***: p<0.005, ns: not significant). **C**. Corrected total fluorescence data was stratified based on the patient immune status for AK (n=5 IC, n=4 IS), BD (n=3 IC, n=5 IS), and cSCC (n=6 IC, n=5 IS). Statistical analysis was performed with t-test.

### PRKDC inhibition reduced AKT phosphorylation, proliferation, and motility in cSCC lines

We tested PRKDC inhibitors in our cell lines. We used NU7026 (NU), which shows a high specificity to PRKDC (Peddi et al., 2010). NU treatment reduced PRKDC phosphorylation in Both pM1, and in Met1 only this was associated with a concomitant decrease in pAKT (Figure 4A). Although PRKDC activation is related to DNA repair following damage, neither Met1, nor Met2 showed any significant difference in γH2Ax foci per cell when treated with NU. pM1 cells on the other hand showed a reduction of foci when treated with NU. (Supplementary Figure S2A and B). This suggested that while control of DNA damage may be important in premalignant lines, in cSCC, PRKDC inhibition over the timeframes we used does not exacerbate DNA damage, suggesting that PRKDC functions in some another capacity in these lines. NU treatment did not reduce viability in the cSCC lines at IC50 doses (Figure 4B), and immunocompetent lines showed similar responses (Supplementary Figure S3A and B). Whilst no viability loss was seen in cSCC at lower doses of NU, 5uM NU did cause a significant reduction in the size of pM1 and Met2 colonies over a 10-day time course (Figure 4C and D), suggesting PRKDC inhibition may drive pM1 and Met1 cells towards differentiation. Met1 cells did not typically grow in colonies, limiting their suitability in this assay (Supplementary Figure S3C). However, migration of Met1 and Met2 cells across a scratch wound did significantly decrease with NU treatment (Figure 4E and F and Supplementary Figure S3D). Met1 cells grown on de-epidermalised dermis in 3D organotypic culture showed reduced expression of the proliferation marker Ki67 (Figure 4G). LTURM34 (LTURM), an alternative PRKDC inhibitor (Morrison et al., 2016), had no measurable effect on either IC or IS cSCC lines. As with NU, the IS and IC1 cells showed no significant response (Supplementary figure S3E and F). Together, our results indicate that whilst PRKDC inhibition does not directly impact cSCC viability, it affects both proliferation and migration of the cSCC keratinocytes.

**Figure 4:**
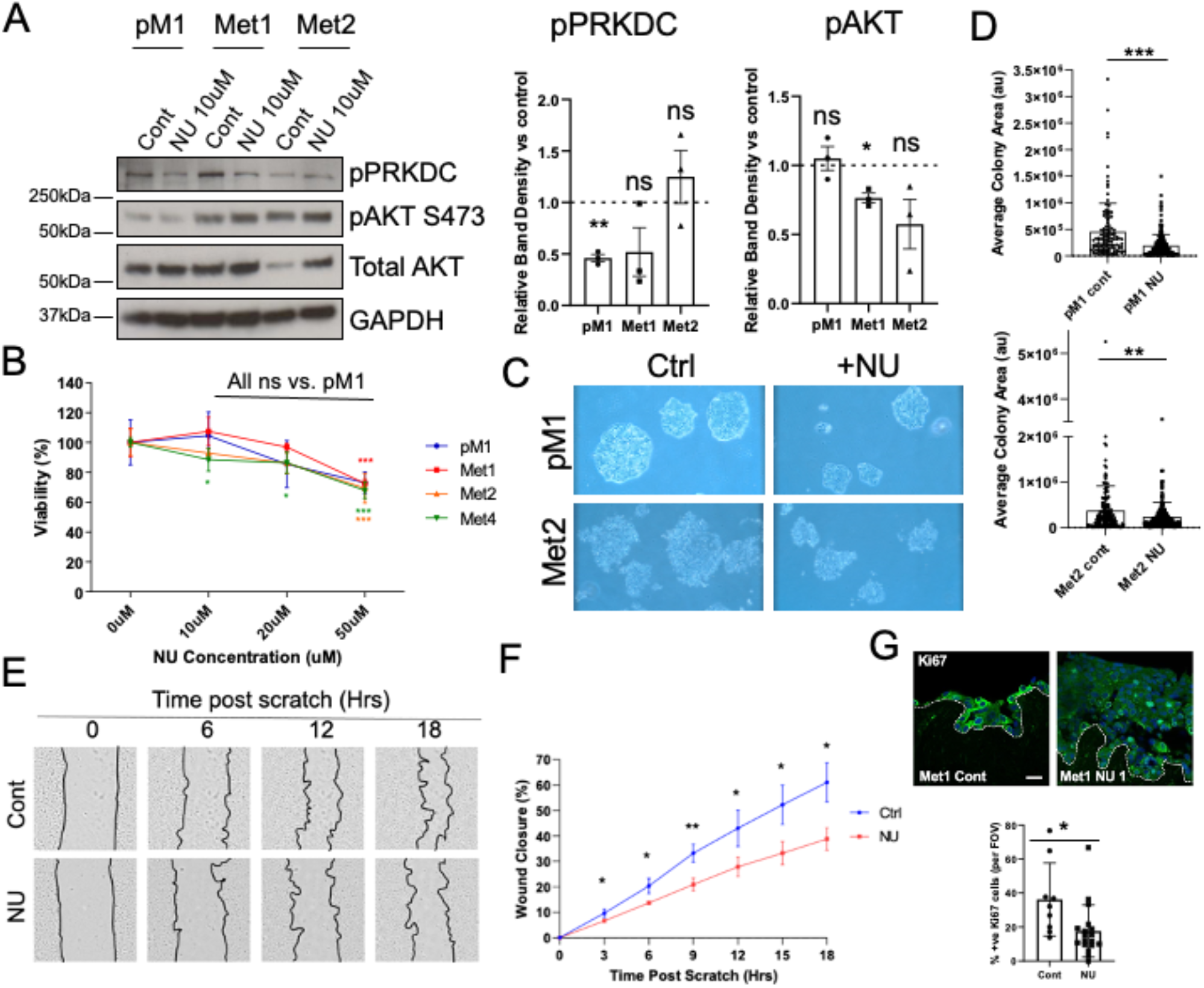
PRKDC inhibitor NU7026 reduces cSCC colony formation and migration across a scratch wound. **A**. Cell lines representing premalignant (pM1), initial cSCC (Met1) and recurrent cSCC (Met2) were treated with vehicle or NU7026 (NU) for 20hrs, Cells were harvested, and lysates analysed by immunoblot by probing as indicated, Accompanying quantification of the mean relative band density of NU treatment vs the matching control is shown for pPRKDC and pAKT in each cell line (±SEM). GAPDH was used for loading control correction for pPRKDC, and total AKT for pAKT. Statistical analysis was performed by t-test for each cell line control vs treatment (**: P<0.01, *: P<0.05, ns: not significant, n=3 for each). **B** cSCC cell lines were treated with NU7026 at the indicated concentrations for 24hrs, then viability was measured with MTT assay. Data are shown as mean ± SD (n=4 per cell line per condition). Statistical analysis was performed with one way ANOVA with Tukey’s Post Test. **C**. pM1 and Met2 lines were seeded at low density and treated with vehicle or 5uM NU. Media with fresh drug was refreshed every other day. At day 10 post seeding, images were taken by light microscopy, with representative examples shown. The average colony area was measured across a n=3 for each cell line condition, with statistical analysis performed by t-test (***: P<0.005). Met1 cells less readily grow in colonies, limiting their use for this analysis (included in Fig S). **D**. Mitomycin-C treated Met1 cells were pre-treated with 5uM NU for 3hrs, and then scratched. Wound healing was measured overnight up to 15 hours. Representative images at 3-hour timepoints are shown with accompanying graphs of % wound healing (n=3). Statistical analysis was performed by t-test for each timepoint. **E**. Organotypic cultures of Met1 cells were established on de-epidermalised dermis and treated with vehicle or 5uM NU. 10 days after rising to the air-liquid interface cultures were fixed, stained with Ki67 antibody, and imaged with confocal microscopy. Representative images are shown. The % of +ve cells per FOV were quantified, with the mean ±sd shown. Statistical analysis was performed by t-test. Scale bar: 50um.

### PRKDC inhibition increases differentiation marker keratin 10 in premalignant cells, and alters YAP expression and localization

NU treatment increased the differentiation marker keratin 10 in pM1 cells (Figure 5A). Previous work has also suggested a role for PRKDC in keratinocyte differentiation (Molinuevo *et al*., 2020). Therefore, we hypothesized that inhibition of PRKDC could promote terminal differentiation in our cSCC lines. Upon NU treatment we were able to detect keratin 10 positive Met1 cells in organotypic culture (Figure 5B). We sought to determine potential underlying mechanisms for this subtle increase in keratin 10. Both YAP1 localization and expression influence keratinocyte differentiation (Walko et al., 2017; Yuan *et al*., 2020), so we explored this within our cSCC lines. Consistent with its known roles in cancer biology, overexpression of nuclear YAP1 leads to SCC formation in mice (Vincent-Mistiaen et al., 2018), total YAP1 expression was higher in Met1 and Met2 vs pM1 (Figure 5C and D). NU treatment decreased YAP1 expression in Met1 and Met2, whilst pM1 showed no sensitivity (Figure 5D). Next, we performed immunostaining on NU treated pM1 and Met1 cells. Nuclear YAP1 was reduced in NU treated pM1 cells and total YAP1 was reduced in NU treated Met1 cells (Figure 5E and F). These changes were confirmed by sub-cellular fractionation (Figure 5G). Together, our findings suggest that PRKDC inhibition with NU not only induces keratinocyte differentiation but also controls YAP1 shuttling from the nucleus to the cytoplasm.

**Figure 5:**
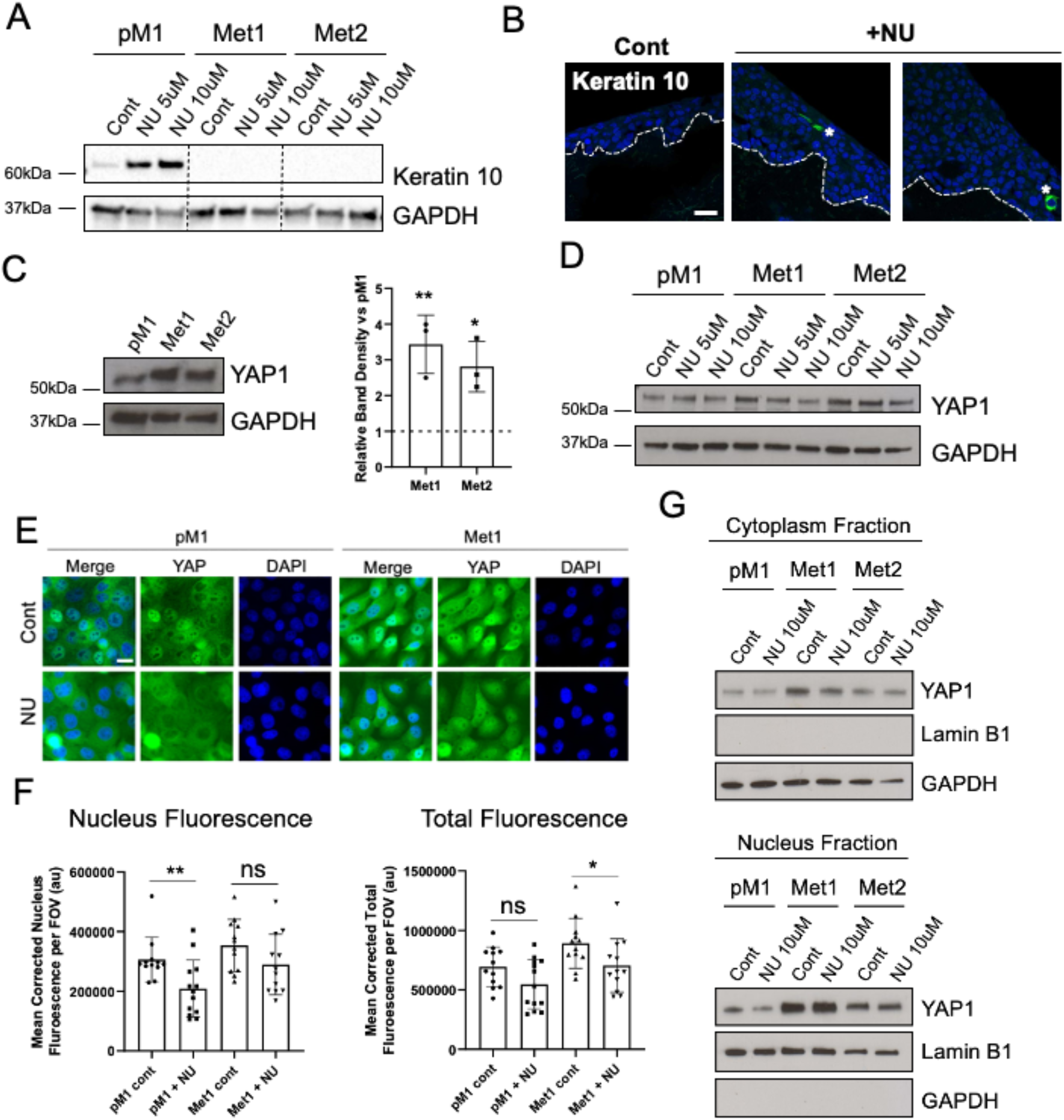
PRKDC inhibitor NU7026 induces differentiation in premalignant cells and changes YAP1 expression and localisation. **A**. Cell lines representing premalignant (pM1), initial cSCC (Met1) and recurrent cSCC (Met2) were treated with vehicle or NU7026 (NU) as indicated for 20hrs, Cells were harvested and lysates analysed by immunoblot by probing as indicated, **B** Organotypic cultures of Met1 cells were established on de-epidermalised dermis and treated with vehicle or 5uM NU. 10 days after rising to the air-liquid interface cultures were fixed, stained with Keratin 10 antibody, and imaged with confocal microscopy. Representative images are shown. * denotes positive cells for keratin 10. Scale bar: 50um. **C**. pM1, Met1 and Met2 lines were lysed and subjected to immunoblot and probed as indicated. Relative band density was then calculated for YAP1, corrected to the GAPDH loading control. Data are presented as mean ±sd of Met1 and Met2 vs pM1 (set to 1 and represented by dotted line). Statistical analysis was performed by ANOVA with Tukey’s post-test (n=3). **: P<0.01, *: P<0.05. **D**. The same lines were treated with vehicle or NU as indicated for 20hrs, and then lysed and subjected to immunoblot. **E**. pM1 and Met1 cells were treated with vehicle or 10uM NU for 20hrs, then fixed and imaged with fluorescence microscopy, with representative images shown. **F**. Graphs of corrected total fluorescence was measured both for the total cell, and cell nucleus only, with mean data ±sd shown. Statistical analysis was performed by t-test with Welch’s post-test (n=2 experiments, with a minimum of n=6 FOV per condition per experiment). **: P<0.01; *: P<0.05; ns: not significant. Scale bar: 20um. **G**. pM1, Met1 and Met2 cells were treated with vehicle or 10uM NU for 20hrs, and then subjected to subcellular fractionation to separate nucleus and cytoplasmic fractions. Lysates were subjected to immunoblot and probed as indicated. Lamin B1 and GAPDH are used as purity/loading controls for nuclear and cytoplasmic fractions respectively.

### Phosphorylation of AKT substrates NDRG2 and YAP1 are increased in cSCC

By Western blot, AKT substrates were markedly upregulated in the pM1 compared to normal keratinocytes, and Met1, 2 and 4 lines show different patterns of substrate phosphorylation (Figure 6A). These could help inform some of the functions AKT drives in cSCC progression, and represent potential druggable targets that are not associated with the risks of pan-AKT inhibition. We cross-referenced our phosphoproteomic dataset with the PhosphoSitePlus database to identify putative AKT substrates (Supplementary Table ST3). There was a total of 9 AKT substrates that showed a significant increase or decrease in at least one of the three Met lines (Figure 6B and C). Across all three cSCC cell lines, the AKT substrate with the greatest fold change in phosphorylation when compared to pM1 was NDRG2 (at T330 and S332) (Figure 6B). Also, we saw up-regulation of YAP1 phosphorylation at S127,128 and 131. In the Met1 and Met2 cell lines, there was a shift from the cytoplasmic localization of YAP1 in the pM1 cells to a more nuclear distribution in the Met1 cells (Figure 6E). This was consistent with the expression pattern in tissues samples, with a cell surface/cytoplasmic distribution in normal healthy skin, changing to predominantly nuclear distribution in SCC, with both AK and BD showing intermediate expression patterns (Figure 6F). NU treatment significantly reduced nuclear pSerine 127 YAP1 in Met1 cells but not in the pM1 cells (Supplementary figure S4).

**Figure 6:**
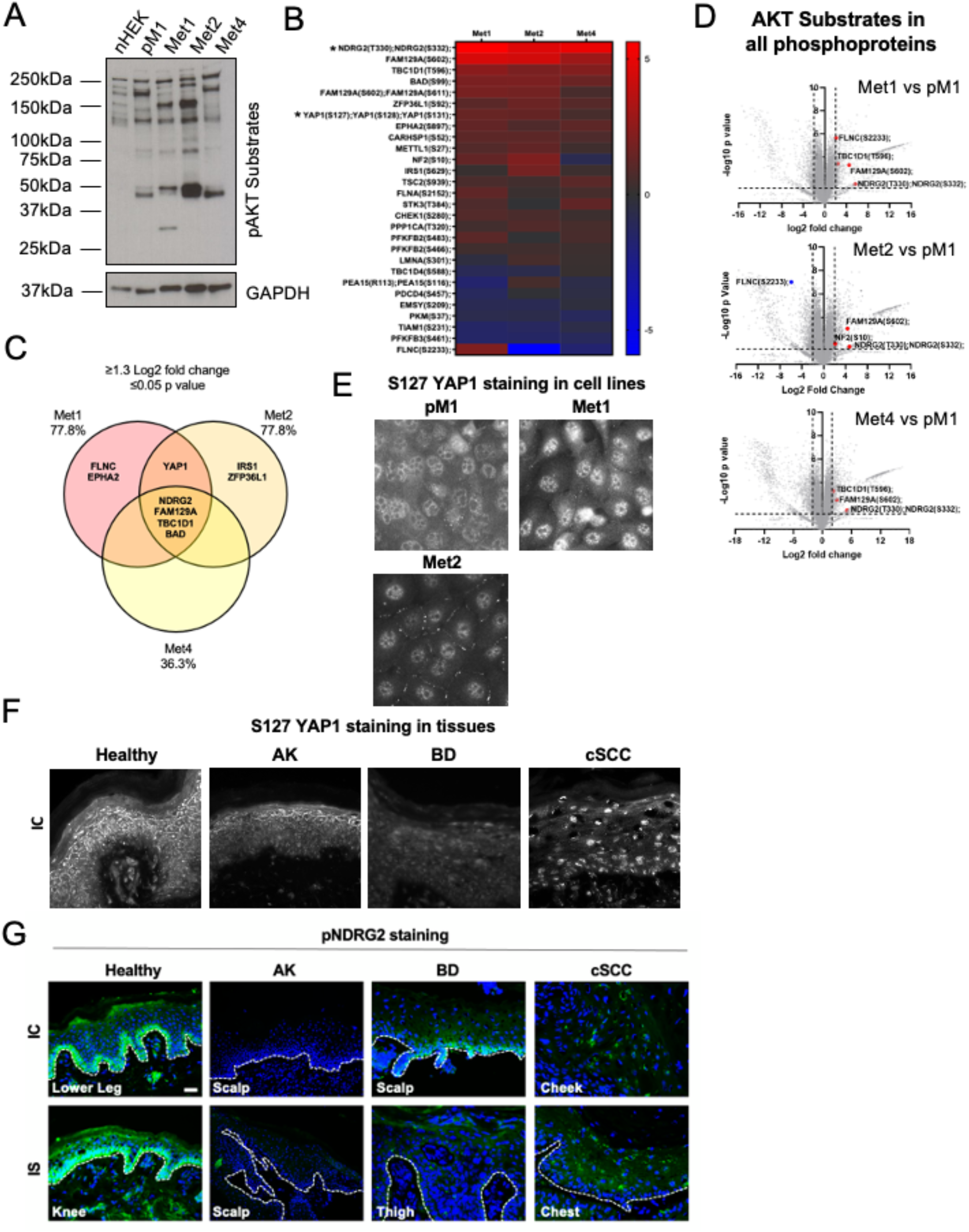
NDRG2 is the most highly phosphorylated AKT substrate in cSCC. **A**. Normal Human Epidermal Keratinocytes (nHEKs), premalignant (pM1), and the cSCC lines Met1, Met2 and Met4 were lysed and analysed by immunoblot with the indicated antibodies. **B**. Heatmap of putative AKT substrates (based on PhosphoSitePlus database) that are significantly altered in at least one of the three Met lines vs. pM1 based on mass spectrometry analysis. **C**. Venn diagram indicates the phosphoproteins upregulated in each of the cell lines compared to pM1 **D**. Volcano plots of all phosphoproteins changed in **(i)** Met1, **(ii)** Met2, **(iii)** Met4, vs. pM1, with AKT substrates with Log2 fold change ≥2, and –Log10 p value ≥1.3 highlighted in each plot. **E**. Representative expression of pS127 YAP1 in pM1, Met1 and Met2 lines. Graph shows the change in nuclear/cytoplasmic ratio in the lines. **F** and **G**. Frozen tissues representing Healthy, Actinic Keratosis (AK), Bowen’s Disease (BD), and cSCC were stained for pS127 YAP1 (F) and pNDRG2 (G), with representative epifluorescence images shown for both immunocompetent (IC) and immunosuppressed (IS) patients. Scale bars: 20um

We explored the role of NDRG2 further in cSCC. Studies have implicated NDRG2 in a variety of different cancers, including, but not limited to lung, pancreatic, glioblastoma (Hu et al., 2016), where generally, NDRG2 expression of NDRG2 is decreased. Pro-apoptotic, anti-proliferative, and anti-migratory roles have been associated with NDRG2. NDRG2 can inhibit AKT via activation of PTEN. Likewise, NDRG2 depletion results in elevated AKT (Nakahata et al., 2014). In gastric cancer cells, AKT inhibition reduces NDRG2 phosphorlyation, rendering the cells more sensitive to apoptosis (Tao et al., 2013). Staining in oral squamous cell carcinoma shows an inverse relationship between NDRG2 and pAKT, with over-expression of NDRG2 reducing levels of pAKT (Furuta et al., 2010).

Based on these previous reports, we hypothesised that NDRG2 expression and activity in our cSCC cells was likely altered, with the elevation in phosphorylation representing either an inhibitory modification on its normal tumour suppressor activity, or the gain of aberrant pro-tumour functions. We began by assessing expression of pNDRG2 (T330 and S332) in our panel of healthy, AK, BD, and cSCC patient tissues. pNDRG2 was expressed in healthy tissue, limited to the more basal and granular layers of the epidermis, consistent with NDRG2 being an AKT2 substrate (O’Shaughnessy et al 2007a). pNDRG2 expression was low across AK and BD tissues and in SCC (Figure 6G).

Nuclear NDRG2 was significantly decreased in Met1 and Met2 lines compared to pM1 (Supplementary Figure S5A). Phosphorylated NDRG2 was increased in Met1 and Met 2 compared to pM1, with discreet foci in the nuclei of pM1 cells and more widespread nuclear expression in Met1 and 2 (Supplementary Figure S5B).

### Targeting NDRG2 with the iron chelator Dp44mT reduces cSCC viability

The iron chelator Dp44mT (Dp) induces NDRG2 expression in hepatocellular carcinoma cells (Wang et al., 2014), so we explored its effects in cSCC. In Met1 and Met2 cell lines, Dp treatment decreased levels of both NDRG2 and pNDRG2, with no change in corresponding pM1 cells (Figure 7A and B). Total AKT and pAKT was also decreased in Met1 and Met2, whilst EMT proteins e-cadherin and vimentin remained unchanged (Figure 7A). By immunostaining, Dp treatment altered NDRG2 expression in both the pM1 and Met1 lines. Nuclear expression of NDRG2 was reduced with Dp treatment in pM1 cells, while cytoplasmic expression of NDRG2 was increased with Dp treatment in Met1 cells (Supplementary Figure S5C).

**Figure 7.**
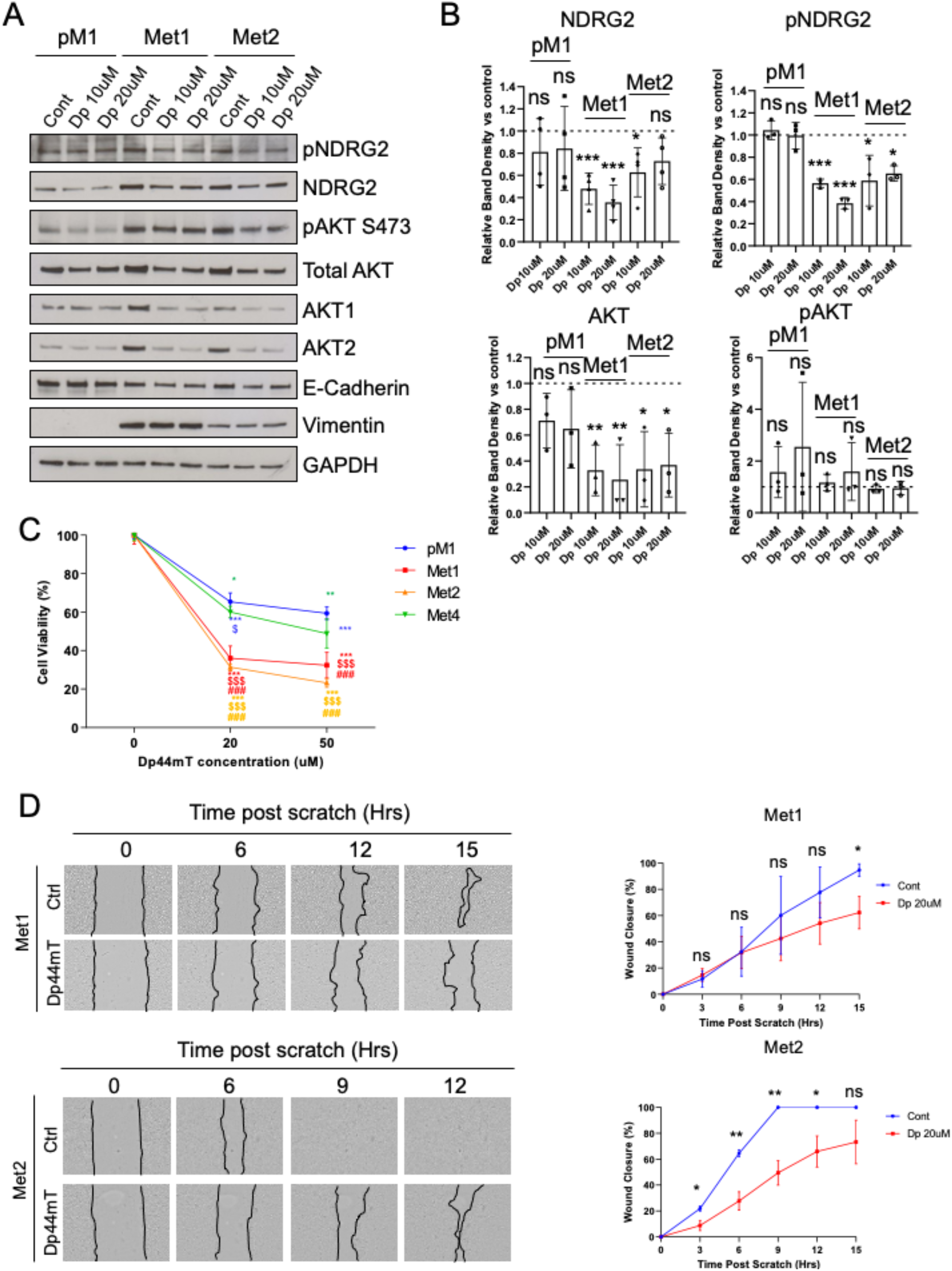

Based on these changes, we were interested to see if Dp treatment changed the viability of the cSCC cell lines. Dp treatment reduced viability in the pM1, Met1, Met2 and Met4 lines, but it was particularly effective in the Met1 and Met2 lines (Figure 7C). In addition, Dp caused a significant reduction in migration across a scratch wound in Met1 cells by 15hrs post-scratch, and at all time points in Met2 cells (Figure 7D). We performed similar experiments in the immunocompetent cSCC lines. Again, Dp treatment reduced both NDRG2 and pNDRG2 levels in IC12 cells by immunoblot. Curiously, this was concomitant with unchanged AKT1 levels and decreased AKT2 levels, and an overall increase in pAKT (Supplementary figure S6A). Importantly, Dp treatment still resulted in viability loss in all three immunocompetent cSCC lines (Supplementary Figure S6B).

### Understanding the overall landscape of AKT signalling in SCC cell lines uncovers therapeutic vulnerabilities

Our comprehensive analysis of phosphosubstrates, and through KSEA analysis, highly activated kinases, provides an overall picture of the control of AKT phosphorylation and downstream protein targets throughout SCC progression in the isogenic lines. Not only do PI3 kinase and mTOR act to phosphorylate AKT, in our tumour lines, PRKDC is also an important kinase controlling AKT phosphorylation. AKT then phosphorylates among other targets, NDRG2, which controls cSCC cell viability. Together, these data suggest increased pNDRG2 drives both cSCC viability and motility, and strategies to inhibit this signalling axis could represent a promising therapeutic approach.

## Discussion

In skin there is a complex relationship between AKT1 expression in the upper layers of the epidermis and AKT2 in the lower epidermis (O’Shaughnessy et al., 2007a and b). While AKT1 activation seems to protect against the effects of UV (Sully et al., 2013), and is required for strong epidermal barrier function (Janes et al., 2003; Thrash et al., 2006; Naeem et al., 2015; Rogerson et al., 2021), AKT2 activation is associated with cSCC progression (O’Shaughnessy et al., 2007b) and proliferation during tumour progression and wound healing (Pankow et al., 2006). Understanding the contribution of AKT1 and AKT2 in SCC by tracking upstream kinases and downstream up- and down-regulated targets will allow more targeted drug interventions to be used to control the output of AKT signalling in cSCC. We show a specific upregulation of PRKDC function during tumour progression and increased phosphorylation of both NDRG2 and YAP1 during tumour progression. We confirm that PRKDC inhibition drives cSCC keratinocytes towards terminal differentiation and that inhibiting NDRG2 by iron chelation reduces cSCC viablility. We demonstrate for the first time in cSCC that phosphoproteomics is a powerful approach to identify key kinases in cSCC progression that can be effectively targeted with small molecule inhibitors.

cSCCs do not undergo normal keratinocyte differentiation. Strategies that can restore normal differentiation, often referred to as ‘Differentiation Therapies’, are an attractive option in delaying or preventing tumour progression (Hugues de The, 2017). Our findings suggest that PRKDC inhibition with NU may induce keratinocyte differentiation by shuttling YAP1 from the nucleus to the cytoplasm. There are few data on the link between YAP1 localisation and DNA damage in keratinocytes, apart from Jun Kinase phosphorylating YAP1 to regulate apoptosis (Tomlinson et al., 2010). AKT phosphorylation of YAP1 at Serine 127 is known to control localization to the cytoplasm, and in so doing protecting against p73 mediated apoptosis (Basu et al., 2003). Paradoxically, we see nuclear YAP1 increase in tumour progression both in cell lines *in vitro* and in our tumour samples (Debaugnies et al., 2018; Vincent-Mistiaen et al., 2018), suggesting that the roles of YAP1 in cSCC are distinct to those in normal homeostasis. Understanding the differential response of YAP1 to AKT in normal keratinocytes, premalignant keratinocytes and during tumour progression would potentially allow for effective targeting of this protein in cSCC (Walko et al., 2017; Jia et al., 2016)

DNA damage induces keratinocyte terminal differentiation as a protective mechanism (Kato et al., 2021). Inhibition of the DNA damage response or sensing of the DNA response prevents the transition from proliferating to differentiating keratinocytes (Mulinuevo et al., 2020). In our cSCC cell lines, inhibition of the PRKDC led to a subtle increase in terminal differentiation. However, this was not linked to an increase in DNA damage that would typically be observed with a reduction of DNA damage sensing and repair. This suggests that there are other, non-DNA damage related functions of PRKDC in keratinocytes. Beyond the immune function of PRKDC evident in *scid* mice (Blunt et al., 1996), clues towards of the function of PRKDC can be gleaned from the expression in normal skin, where the phosphorylated form is confined the to the uppermost layers of the epidermis. This could be attributed to the DNA damage accrued by nuclei during the final stages of epidermal terminal differentiation (Mulinuevo et al., 2020), or here could be a direct role for PRKDC during epidermal terminal differentiation.

Both PI3 kinase and PRKDC therefore contribute to AKT phosphorylation in cSCC. One obvious limitation of these analyses was the inability to detect PI3 phosphorylation. However, treatment with the PI3K inhibitor LY294002 and specific inhibition of the p110a subunit reduces viability of both premalignant cells and the Met1 cell line (Mannella et al., 2021). This suggests that the AKT phosphorylation via PI3K is important in both normal homeostasis and tumourigenesis, while the PRKDC mediated phosphorylation of AKT is important in tumour progression, as no effects on viability were seen in premalignant lines when treated with NU. One potential explanation is a differential effect on AKT1 and AKT2, with NU specifically reducing AKT2 phosphorlyation. Interestingly we saw no effect with another PRKDC inhibitor LTURM (Chandra et al., 2021). We cannot discount the possibility of inhibition of PI3K by NU7026 as being the reason for the difference, however as we didn’t see any effect in premalignant lines, this suggests that NU7026 is specifically inhibiting PRKDC in our experiments.

We saw differential responses in different cell lines, especially in the immunocompetent line under test. Although there was reduced viability, overall AKT phosphorylation was increased. This unexpected elevation of AKT could either represent the homeostatic AKT1 isoform, or the possibility of differing signaling responses in IC vs IS cSCCs. For instance, NDRG2 depletion has previously been shown to elevate AKT by virtue of interactions with PTEN (Nakahata et al., 2014). This suggests that either the increase of AKT1 or maintenance of AKT1 activity and the loss of AKT2 expression and activity induced by Dp treatment was potentially cancer protective. This is consistent with known responses to UV light, HPV infection and cSCC progression, where the loss of AKT1 activity typically presages the increase in overall AKT2 activity (O’Shaughnessy et al., 2007b; Sully et al., 2013). Understanding ways to activate AKT1 activity while maintaining inhibition of AKT2 will be important in the treatment of cSCC. Rapamycin treatment can activate AKT1 specifically in keratinocytes (Sully et al., 2013), and use of rapamycin reduces cSCC incidence after transplant, and mTOR inhibition by everolimus is used to treat bladder cancer (Hoogendijk-van den Akker et al., 2013)

Little is known about NDRG2 function in the epidermis. Our expression data in normal skin suggest that phosphorylated NDRG2 (T330/T332) is localized to the basal layer and the uppermost layers, which correlates well with the expression of active AKT1 and AKT2 respectively (O’Shaughnessy et al., 2007a). The loss of pNDRG2 in SCC suggest that phosphorylation at these sites promotes degradation of NDRG2. NDRG2 loss has been reported in a range of cancers (Gu et al., 2020; Tamura et al., 2017). Conversely, NDRG2 overexpression reduces cSCC growth (Wang et al., 2015). NDRG2 regulates adherens junctions in the colon and reduced NDRG2 leads to reduced expression of E-Cadherin (Wei et al., 2020) which we also observe upon Dp treatment, which also reduced NDRG2 expression in cSCC keratinocytes. Loss of E-cadherin prevents collective migration of keratinocytes (Tu et al., 2019), and consistent with this we see reduced migration across a scratch wound. However, loss of E-cadherin also correlates with increased invasion due to epithelial-mesenchymal transition (EMT). It would be important to determine whether markers of EMT are being increased in expression in response to iron chelator treatment in cSCC, although in other tumours iron chelator treatment reduces metastasis (Wang et al., 2014). Together, our findings suggest increased AKT-mediated phosphorylation of NDRG2 helps drive cSCC viability and motility, and strategies to inhibit this axis could represent a promising therapeutic angle.

In summary, we demonstrate that phosphoproteomics represents a powerful approach to identifying key kinases in cSCC progression that can be effectively targeted with small molecule inhibitors. These data have significant implications for development of effective targeted therapies, a current area of major unmet clinical need in the treatment of advanced cSCC.

## Materials and Methods

### Antibodies and Reagents

#### Rabbit Polyclonal Antibodies

Keratin 10 (BioLegend Poly19054) (1:1,000 WB; 1:50 IF); Keratin 5 (BioLegend Poly19055) (1:1,000 WB); Phospho-AKT (Ser473) (CST #9271) (1:1,000 WB; 1:30 IF); Total AKT (CST #9272) (1:1,000 WB); Phospho-(Ser/Thr) Akt Substrate Antibody (CST #9611) (1:1,000 WB); Phospho-DNA-PK (Thr2609) (ThermoFisher #PA1-29541) (1:1,000 WB; 1:50 IF); Ki67 (GeneTex #GTX20833) (1:50 IF); Phospho-Histone H2A.X (Ser139) (CST #2577) (1:200 IF); Lamin-B1 (Abcam #ab16048) (1:1,000 WB); NDRG2 (CST #5667) (1:1,000 WB, 1:50 IF).

#### Rabbit Monoclonal Antibodies

E-cadherin (CST #3195) (1:1,000 WB); AKT1 (CST #2938) (1:1,000 WB); AKT2 (CST #3063) (1:1,000 WB); YAP (CST #14074) (1:1,000 WB; 1:50 IF)

#### Mouse Monoclonal Antibodies

Vimentin (CST #3390) (1:1,000 WB); GAPDH (Millipore Sigma) #MAB374 (1:2,000 WB)

#### Sheep Polyclonal Antibodies

Phospho-NDRG2 (Thr330+Ser332) (MRC Protein Phosphorylation & Ubiquitylation Unit, Dundee #S971A) (0.5µg WB; 5µg Cell IF; 10µg Tissue IF).

#### Secondary Antibodies

##### Western blot

Swine anti-rabbit immunoglobulins/HRPs (Dako #P0399) (used 1:500-1:5,000); Goat anti-mouse immunoglobulins/HRPs (Dako #P0447) (used 1:500-1:5,000); Rabbit anti-sheep immunoglobulins HRPs (Dako #P0449)

##### Immunofluoresence

Goat anti-rabbit IgG Cross-adsorbed secondary antibody Alexa Fluor 488 (Invitrogen #A-11008) (1:700), Goat anti-rabbit IgG Cross-adsorbed secondary antibody Alexa Fluor 568 (Invitrogen #A-11011) (1:700); Donkey anti-sheep IgG Cross-adsorbed secondary antibody Alexa Fluor 488 (Invitrogen #A-11015) (1:700).

##### Inhibitors

NU7026 (#N1537) and Dp44mT (#SML0186) were products of Merck. LTURM34 (#S8427) was from Selleckchem. Mitomycin C (Thermo Fisher J63193.MA)

### Cell lines and culture

The cell line series of pM1 (pre-malignant, forehead), Met1 (initial cSCC, dorsal hand), Met2 (cSCC recurrence of Met1), and Met4 (lymph node metastasis of the tumour) are as previously described (Proby et al., 2000). All four lines originate from the same patient, a 45-year-old immunosuppressed male. IC1 SCC (right temple) and IC1 Met (right preauricular lymph node metastasis of IC1 SCC) originated from a 77-year-old immunocompetent male. IC12 cells are from the left calf of an 87-year-old immunocompetent female. Normal human epidermal keratinocytes were obtained from Merck. All lines were maintained in 3:1 DMEM (Gibco, 11960044): F12 media (Gibco, 11765054), with 10% FBS, 5% pen/strep, 5% l-glutamine, and 5% RM^+^ supplement (consisting of 5ug/ml Transferrin (Gibco, #11508846), 0.4ug/ml Hydrocortisone (Sigma, #H4881), 1×10^−8^M Cholera Toxin (Sigma, #C8052), 10ng/ml mouse EGF (Gibco, #PMG8043), 5ug/ml Insulin (Sigma, #I5500), 2×10^−11^M Liothyronine (Sigma, #T6397)). Cells were detached for passaging and plating with 0.05% Trypsin-EDTA (Gibco, #25300054).

### Mass Spectrometry Phosphoprotemic Analysis

Phosphoproteomics experiments were performed using mass spectrometry as reported (Rajeeve et al., 2014 Mol Cell Proteomics; Casado et al., 2013 Sci Signaling). In brief, cell pellets were lysed in 8M urea buffer supplemented with phosphatase inhibitors (10 mM Na_3_VO_4_, 100 mM β-glycerol phosphate and 25 mM Na_2_H_2_P_2_O_7_ (Sigma)). 250µg of protein lysate was used for each cell line. Proteins were digested into peptides using trypsin as previously described (Alcolea *et al*., 2012 Mol Cell Proteomics; Montoya *et al*., 2011 Methods). Phosphopeptides were enriched from total peptides by TiO_2_ chromatography essentially as reported previously (Larsen *et al*., 2005 Mol Cell Proteomics). Dried phosphopeptides were dissolved in 0.1% TFA and analysed by nanoflow ultimate 3000 RSL nano instrument was coupled on-line to a Q Exactive plus mass spectrometer (Thermo Fisher Scientific). Gradient elution was from 3% to 35% buffer B in 90 min at a flow rate 250 nL/min with buffer A being used to balance the mobile phase (buffer A was 0.1% formic acid in water and B was 0.1% formic acid in acetonitrile). The spray voltage was 1.95 kV and the capillary temperature was set to 255 °C. The Q-Exactive plus was operated in data dependent mode with one survey MS scan followed by 15 MS/MS scans. The full scans were acquired in the mass analyser at 375-1500m/z with the resolution of 70 000, and the MS/MS scans were obtained with a resolution of 17 500.

MS raw files were converted into Mascot Generic Format using Mascot Distiller (version 2.7.1) and searched against the SwissProt database (December 2018) restricted to Human entries using the Mascot search daemon (version 2.6.0). Allowed mass windows were 10 ppm and 25 mmu for parent and fragment mass to charge values, respectively. Variable modifications included in searches were oxidation of methionine, pyro-glu (N-term) and phosphorylation of serine, threonine, and tyrosine. Peptides with an expectation value<0.05 were considered for further analysis. Phosphopeptides were quantified using Pescal Software (Cutillas & Vanhaesebroeck 2007 Mol Cell Proteomics), which was used to construct extracted ion chromatograms for all the identified peptides across all conditions. The resulting quantitative data was parsed into excel files for further normalisation and statistical analysis.

### Gene Ontology (GO) Analysis

Phosphoprotein lists from each cell line were thresholded based on a fold change vs pM1 of +/- 2. Proteins were analysed using the PANTHER GO tool for Biological Process from The Gene Ontology Resource (Ashburner *et al*., 2000 Nature Genetics; The Gene Ontology Consortium 2021 Nucleic Acids Research; Mi *et al*., 2019 Nucleic Acids Res). Terms were selected with P value <0.05 and false discovery rate (FDR) <0.05. We present the 15 greatest fold change terms that show ≥1.5 fold change in each line.

### Kinase Substrate Enrichment Analysis (KSEA)

KSEA has been described previously (Casado *et al*., 2013 Sci Signal). Our KSEA data was based on Z-score generated using PhosphoSitePlus database. Kinase candidates with a Z-score higher than 1 were used to create a protein-protein interaction network using STRING (Szklarczyk et al., 2021)

### Western Blotting

Cells were lysed in total lysis buffer (20% β-mercaptoethanol, 5% sodium dodecyl sulphate (SDS), 10 mM Tris pH 7.0) and boiled for 10 min at 95 °C. Lysates were separated via 4-20% SDS-polyacrylamide gradient gels (Bio-Rad). Proteins were transferred to nitrocellulose membranes before blocking with 5% milk (Marvel) or bovine serum albumin (Sigma) w/v in PBS-T (0.1% Tween-20 in PBS). Blocked membranes were probed with primary and HRP-linked secondary antibodies diluted in the blocking buffer. Luminol (Santa Cruz Biotechnology) was used to detect protein bands and densitometry was calculated using ImageJ software.

### Human Tissue Samples

Full local ethical approval was granted for use of human tissues from healthy skin (HS), Actinic Keratosis (AK), Bowen’s Disease (BD), and cSCCs (SCC). The source of tissue samples used in this investigation are summarised in Supplementary table ST1. Snap frozen tissues were embedded in OCT and 10µm sections were cut on a cryostat.

### Immunofluorescent Staining

#### Frozen tissues

Tissue sections were defrosted for 15min at RT then fixed in 4% paraformaldehyde (PFA) for 10min. Tissues were then washed 3 times in 1XPBS, and permeabilized with PBS containing 0.1% Triton X-100 for 20min. After a further 3 1XPBS washes, tissues were moved to a humidified chamber and blocked with blocking buffer (PBS with 0.4% gelatin from cold fish and 0.2% Triton X-100) for 45min. Tissues were probed overnight at +4°C with primary antibody diluted in blocking buffer in the humidified chamber. The next day, tissues were washed 3 times in 1XPBS for 5 mins, and then probed with fluorophore-conjugated secondary antibody for 1hr in the dark at RT. After a final 3 1XPBS 5 min washes, coverslips were mounted on tissues using Prolong Gold DAPI solution (Sigma Aldrich). Tissues were left at RT in the dark for mounting to dry, then kept at +4°C for storage.

#### Cell lines

Cells in culture had media removed, a 1XPBS rinse, and then were fixed in 4% PFA containing 0.2% Triton X-100 for 10min. After 3 1XPBS rinses, cells were blocked in blocking buffer (1XPBS with 0.4% fish skin gelatin and 0.2% Triton X-100) for 30min. Primary antibodies diluted in blocking buffer were then added to coverslips for 1hr at RT. After 3 1XPBS washes, fluorophore-conjugated secondary antibody diluted in blocking buffer was added and incubated for 45min at RT in the dark. Covers were given a further 3 1XPBS washes, and then mounted using Prolong Gold DAPI solution. Coverslips were left at RT in the dark for mounting to dry, then kept at +4°C for storage. Imaging was performed on either a Leica DM5000B Epifluoresence microscope or Zeiss confocal microscope. Corrected total cell fluorescence was measured using ImageJ.

### Viability Assays

Cells were plated in a 96 well-plate at ∼2,500 cells/plate. Drug or vehicle (DMSO) were added at the desired concentrations and incubated for 24-48hrs (check figure legends for specific experiments). MTT reagent (Invitrogen, #M6494) was prepared as a 12mM solution in PBS, and 10µl was added to each well. Cells were incubated for 2hrs at 37°C. Media was then removed, and 100µl DMSO added to each well, and pipetted up and down 5 times to resuspend formazan crystals. Plates were then incubated covered from light on a plate shaker for 5min. Absorbance was measured on a plate reader (CLARIOstar, BMG Labtech). Data were calculated based on setting vehicle control as 100% viability, and other treatments as % change from this.

### Colony Assays

5,000 cells were seeded on 10 cm dishes and treated with drug or vehicle (DMSO). Media was changed and drugs replenished every other day. At day 10 post seeding, cells were fixed with 4% PFA, and then images taken on a light microscope. Colony areas were measured on ImageJ.

### Scratch Assays

Cells were grown to full confluence and treated with 5µg/ml Mitomycin-C for 2hrs to stop cell division. Media was removed and cells were washed 3 times with 1XPBS to remove any remaining Mitomycin-C. Cells were incubated with vehicle or drug for 3hrs, and then cells were scratched with a pipette tip. Media was replaced with fresh vehicle/drug, and imaging was performed overnight at 37°C with a CytoSmart Lux2 microscope. Data were calculated based on % of the wound area healed over time.

### Organotypic Culture

pM1 or Met1 cells were seeded on ∼2cmx2cm squares of de-epidermalised dermis at a density of ∼100,000 cells, with vehicle/drugs added as desired. After cells reached confluence, cells were moved to the air-liquid interface. Media and drugs were replenished every other day. After 10 days at the air-liquid interface, cultures were fixed in 4% PFA for 2hrs, and then placed into 70% ethanol. Tissues were then embedded in paraffin. 10µm sections were cut on a microtome. Before staining, tissues were dewaxed (5 min washes in xylene (x2), 100% ethanol (x2), 90% ethanol, 70% ethanol, water). Antigen retrieval was performed using 0.01M sodium citrate buffer pH 6.0 in a microwave, using 2min high heat and 5min med/low heat. Staining then proceeded using the same protocol as for cell lines.

### Nucleus Isolation

Cells were trypsinised and pelleted in a centrifuge, then washed once and spun again in cold 1XPBS. Pellets were resuspended in 100µl ice cold 1XPBS, with the same volume 2Xlysis buffer added (1M Tris/HCl pH7.5, 1M NaCl, 1M MgCl_2_, 10% IGEPAL in MilliQ water). Cells were pipetted 20 times to ensure proper lysis. Lysates were spun at 500xg at 4°C for 30min. Supernatant (cytoplasmic fraction) was collected. Remaining pellets (nucleus fraction) were resuspended and washed once in ice cold resuspension buffer (10mM NaCl, 10mM Tris pH 7.4, 3mM MgCl_2_ in MilliQ water) at 1000xg at 4°C for 5min. All supernatant was removed and then pellets resuspended in an appropriate volume (∼150µl) of resuspension buffer. Both cytoplasmic and nucleus fractions were treated with 2X lysis buffer (Molecular Probes) and boiled at 95°C for 10min. Lysates were then analysed by western blot.

## Acknowledgements

We would like to acknowledge Vinothini Rajeeve for phosphoproteomics analysis, Rob Hearnden for initial data analysis, Rhiannon Laban for providing tissue blocks, Liisa Blowes for assistance with scratch assays and Ros Hannen for assistance with organotypic culture assays. This work was funded by Barts and the London Charity and the British Skin Foundation. RFLO’S,CH and RB conceived the study. RFLO’S, CH and RB designed experiments. RB performed experiments and interpreted data. The manuscript was written by RFLO’S, RB and CH.

## Supplementary Information

**Supplementary figure S1:**
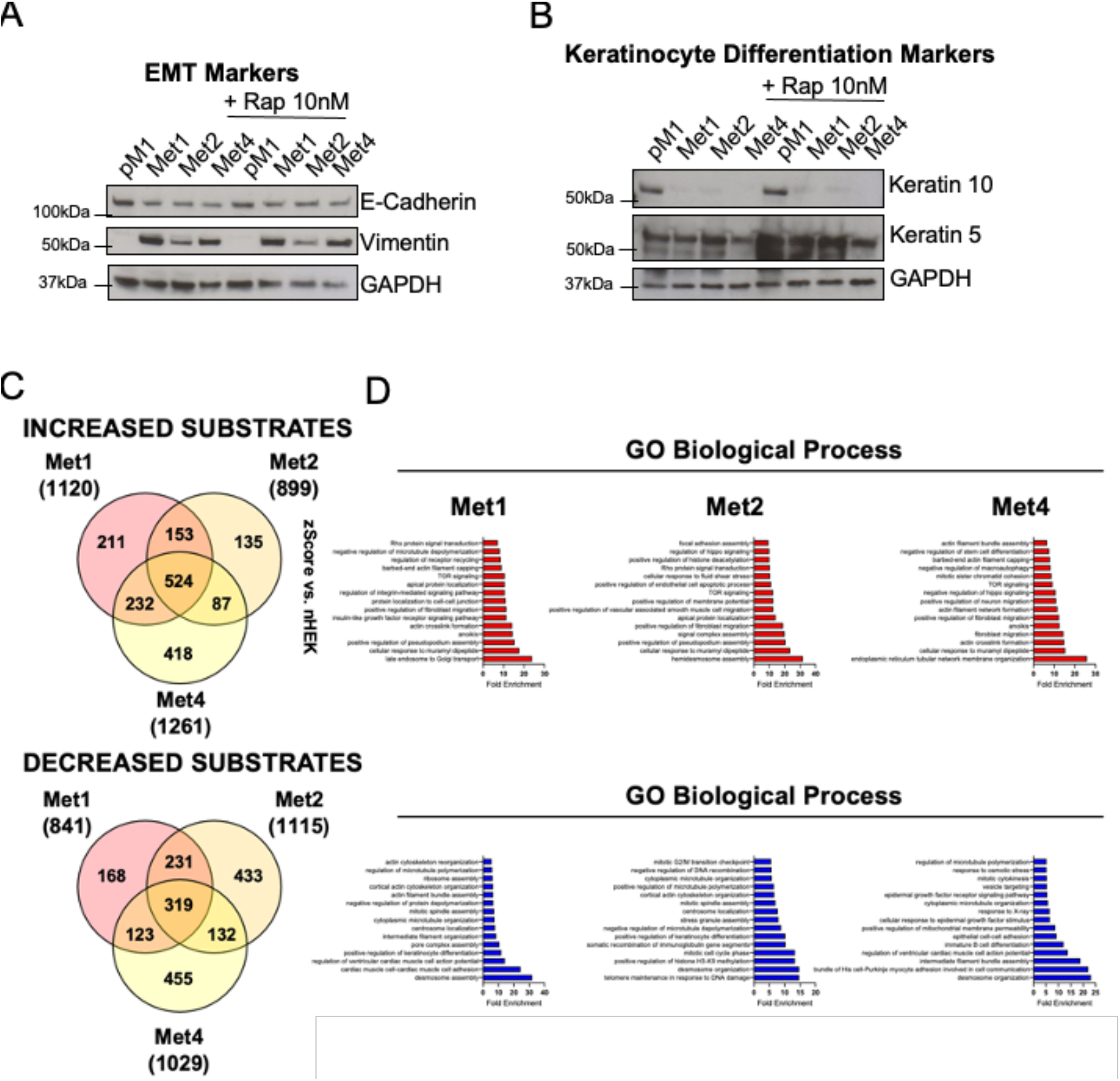
Characterisation of immunosuppressed cSCC cell lines used in study. **A and B**. pM1, Met1, Met2 and Met4 lines were harvested and analysed by Immunoblot either with or without 10nM Rapamycin and probed with markers for EMT **(A)** or keratinocyte differentiation **(B)**. GAPDH was used as loading control. **C**. Venn diagram summary of the total number of phosphorylated residues that are increased in Met1 (initial cSCC), Met2 (recurrence of Met1), and Met4 (lymph node metastasis of Met2), versus a premalignant (pM1) cell line. Thresholds were set to ≥1.5-fold change and p≤ 0.05. **D**. Fold enrichment of the highest 15 Gene Ontology Biological Process terms in each line are shown. Kinases which showed a ≥2 z-score and ≤0.05 p-value in at least one of the three cell lines are displayed.

**Supplementary figure S2:**
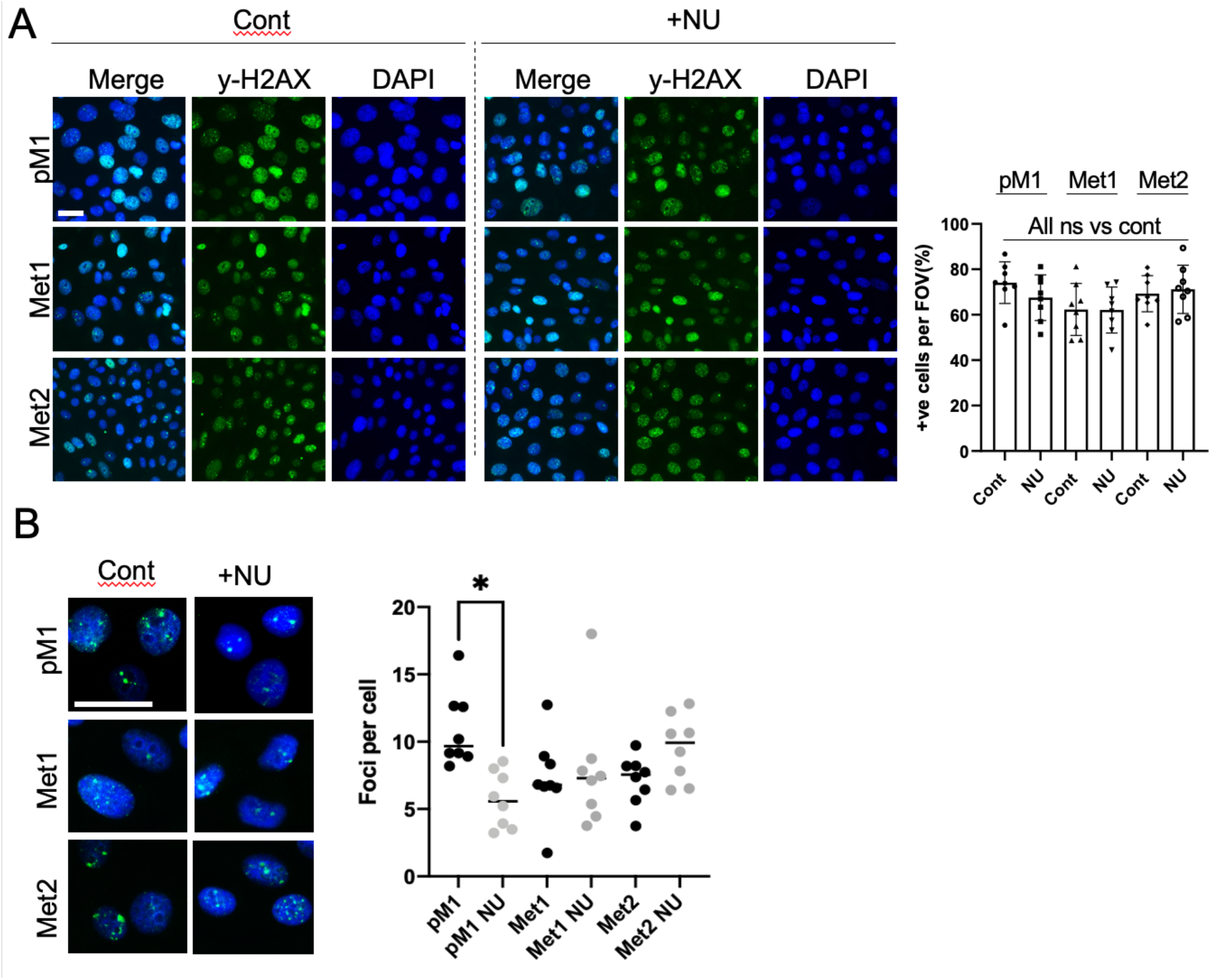
DNA damage marker y-H2AX is not affected by PRKDC inhibitor NU7026 in cSCC. **A and B**. cSCC cell lines were treated with vehicle or 10uM NU7026 (NU) for 24hrs, then fixed and stained with y-H2AX antibody. **A**. Representative fluorescence images are shown. The % cells with y-H2AX puncta per FOV were quantified, with mean values ±sd shown **B**. Counts of foci per cell showed a reduction only in pM1 cells treated with NU. No change was seen in Met1 or Met2 cSCC lines. n=8 FOV in all analyses. P<0.05, ANOVA with Tukey’s multiple comparisons. ns= not significant. Scale bar: 20um

**Supplementary Figure S3:**
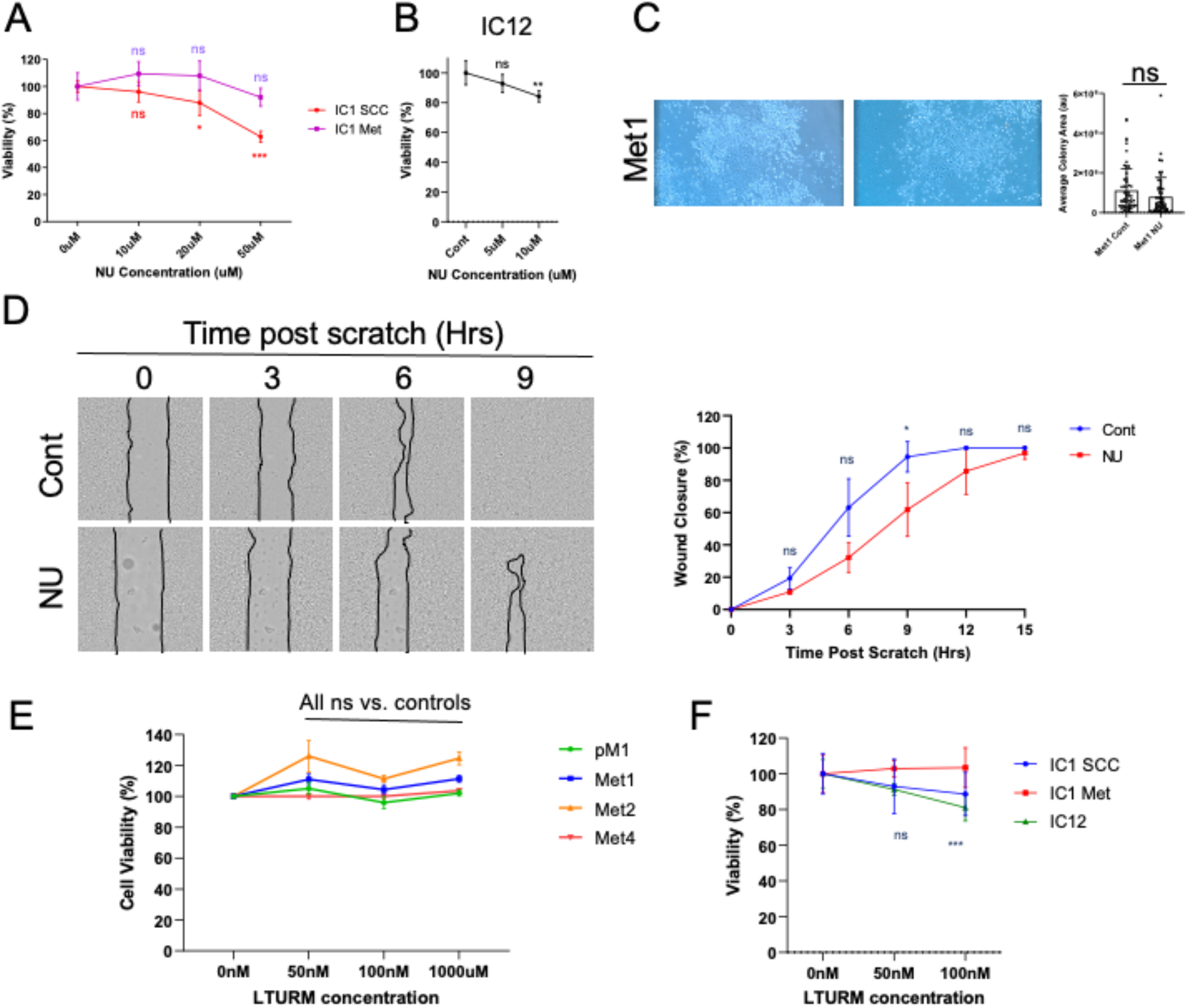
Additional responses of cSCC lines with PRKDC inhibitors. **A**. Immunocompetent cSCC cell lines IC1 SCC and IC1 Met were treated with NU7026 at the indicated concentrations for 48hrs, then viability was measured with MTT assay. Data are shown as mean ± SD (n=4). Statistics was performed with one way ANOVA with Tukey’s Post Test. ***: P<0.05; *: P<0.05. **B**. The IC12 line was treated with NU for 24hrs and analysed as in C (n=4), **: P<0.01. **C**. Met1 cells were seeded at low density and treated with vehicle or 5uM NU. Media with fresh drug was refreshed every other day. At day 10 post seeding, images were taken by light microscopy, with representative examples shown. The average colony area was measured across a n=3 for each cell line condition, with statistical analysis performed by t-test (ns= not significant). **D**. Mitomycin-C treated Met2 cells were pre-treated with 5uM NU for 3hrs, and then scratched. Wound healing was measured overnight up to 15 hours. Representative images at 3-hour timepoints are shown with accompanying graphs of % wound healing (n=3). Statistical analysis was performed by t-test for each timepoint. **E**. pM1, Met1, Met2 and Met4 lines were treated with PRKDC inhibitor LTURM at the indicated concentrations. Data are shown as mean ±SEM (n=4 per line per condition). Statistical analysis was performed as in B. **F**. All three immunocompetent cell lines were treated with LTURM for 24hrs and analysed as before (n=4 per cell line per condition).

**Supplementary figure S4:**
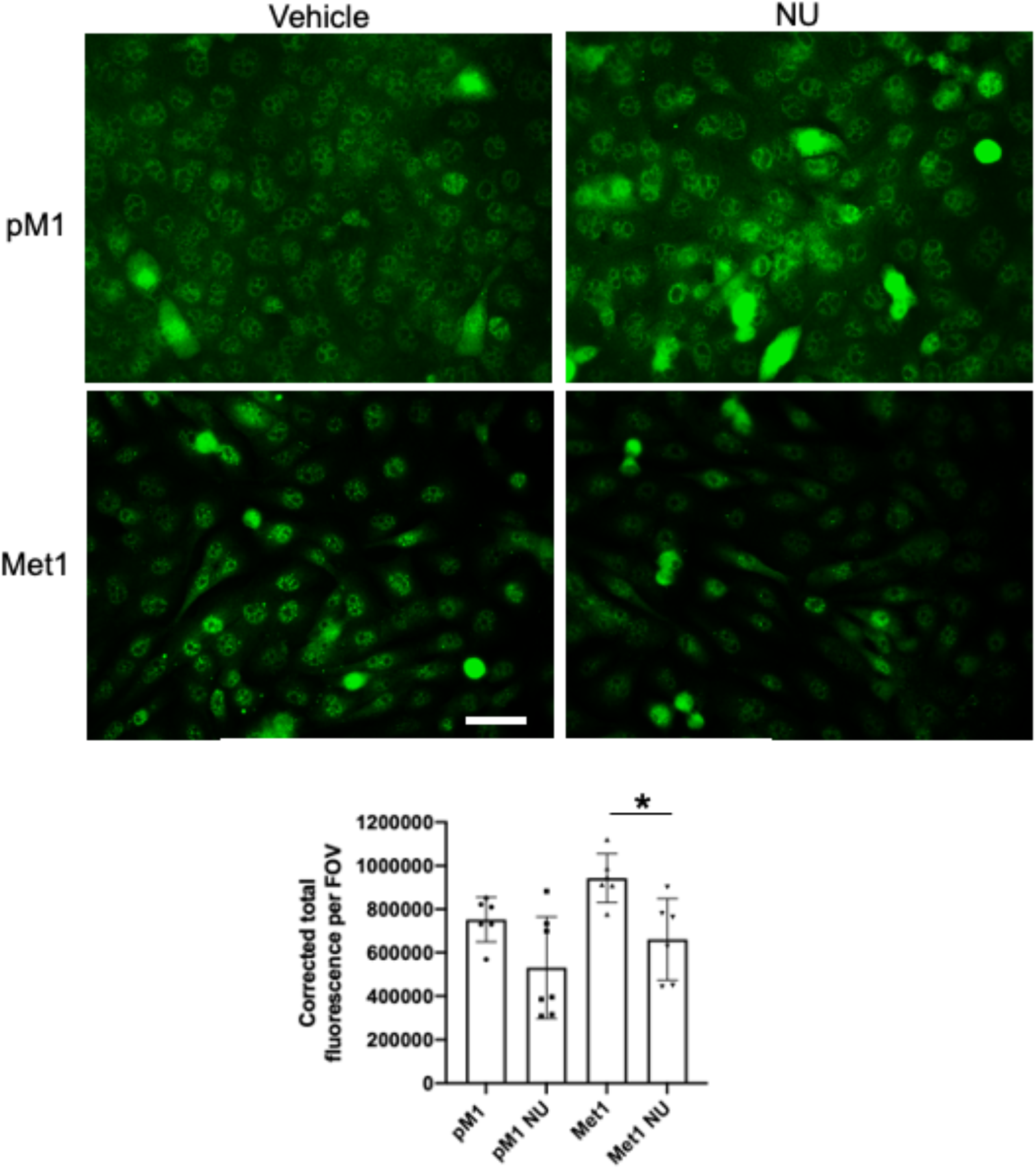
NU treatment reduces nuclear pSer127 YAP1 in SCC cell lines. Representative immunofluorescence of pSer127 YAP1 in pM1 and Met1 cells. *p<0.05 1-way ANOVA followed by post-hoc testing. Bar 30um

**Supplementary Figure S5:**
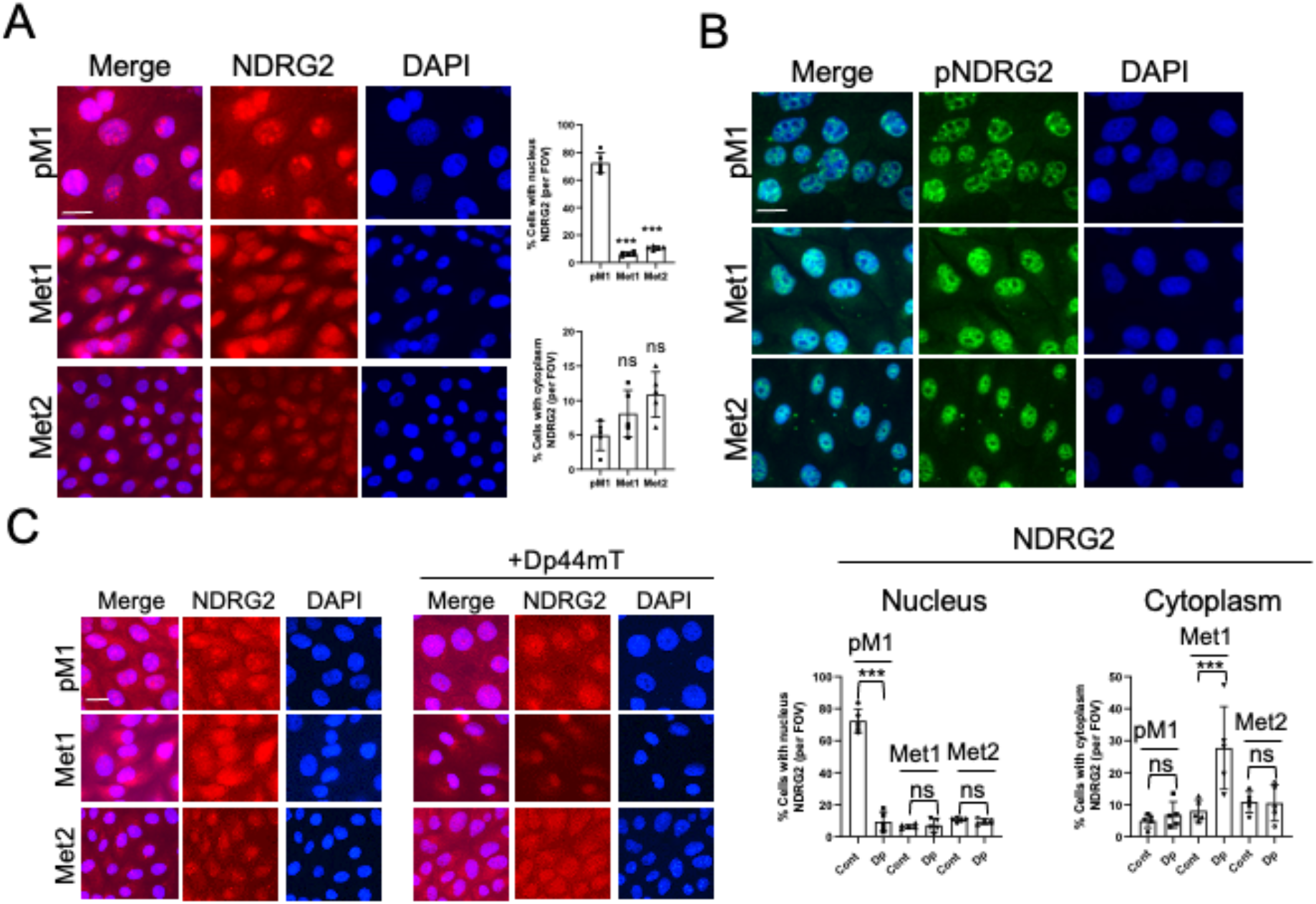
Expression pattern of NDRG2 and response to Dp44mT in cSCC cell lines. **A**. pM1, Met1, and Met2 cells were fixed and stained with NDRG2 antibody. The number of cells per FOV (%) with nucleus or cytoplasmic NDRG2 puncta were quantified, with mean values shown (n=5 FOV). Data were analysed with one way ANOVA and Tukey’s Post Test. ***: P<0.005; ns: not significant. Scale bar: 20um. **B**. The same cell lines were fixed and stained for pNDRG2 antibody. Scale bar: 20um. **C**. cSCC lines were treated with 20uM Dp44mT for 24hrs, then fixed and stained for NDRG2. Analysis was performed as in **A**. Scale bars: 20um.

**Supplementary figure S6:**
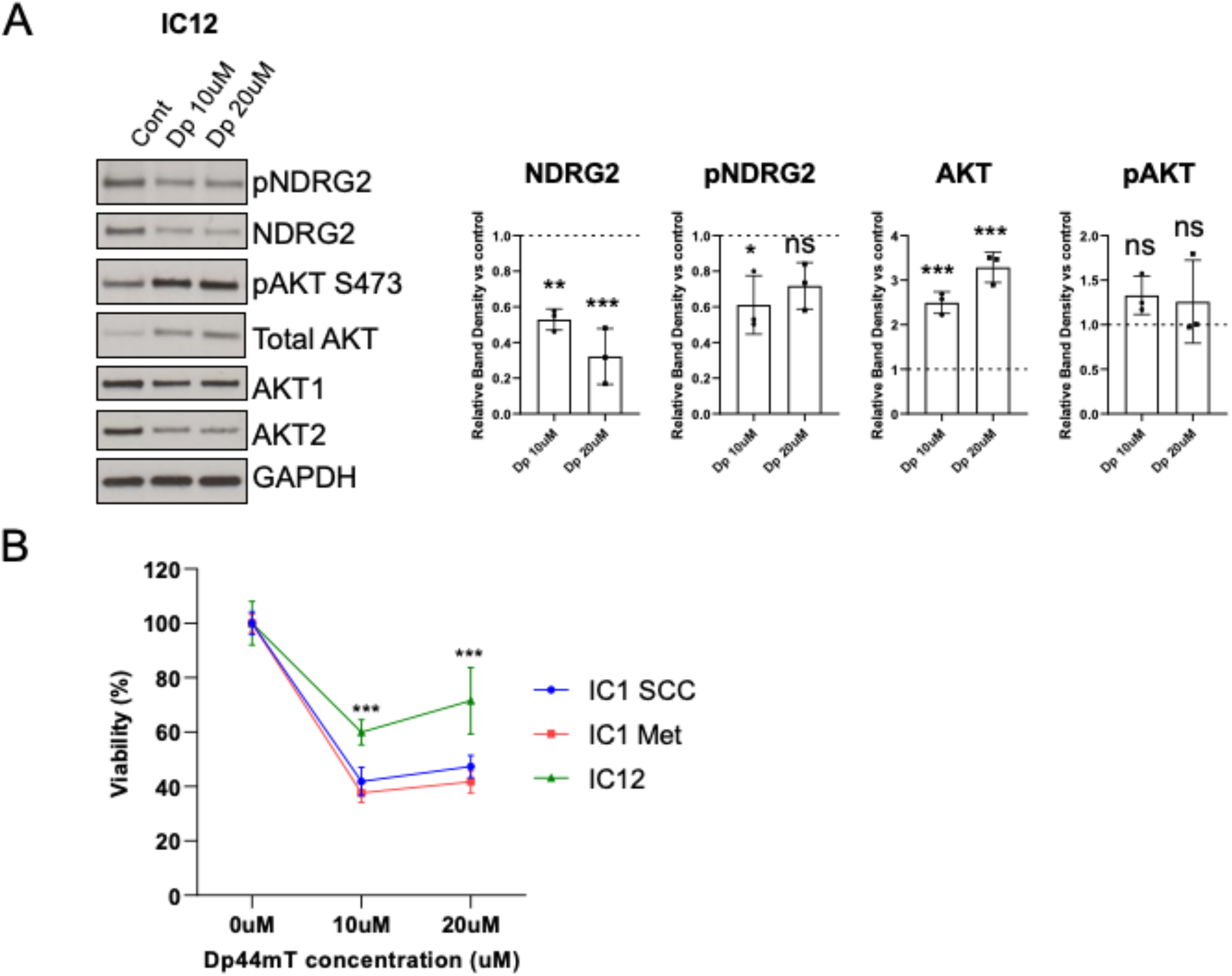
Iron Chelator Dp44mT treatment also reduces NDRG2 and viability in immunocompetent cSCC. **A** Immunocompetent cSCC line IC12 was treated with vehicle or Dp44mT as indicated for 20hrs, and lysates were then harvested and analysed by immunoblot with the indicated antibodies. Quantification of western blot band density is shown, with dotted lines representing the vehicle control for which treatments were corrected to. Statistical analysis was performed by t-test (n=3) (***: P<0.005; **: P<0.01; *: P<0.05; ns: not significant). **B**. Immunocompetent cSCC lines IC1 SCC, IC1 Met and IC12 were treated with Dp44mT as indicated for 24hrs and then viability was measured by MTT assay (n=4 per cell line per condition). Statistical analysis was performed by ANOVA with Tukey’s as a post test.

**Supplementary Table ST1.**
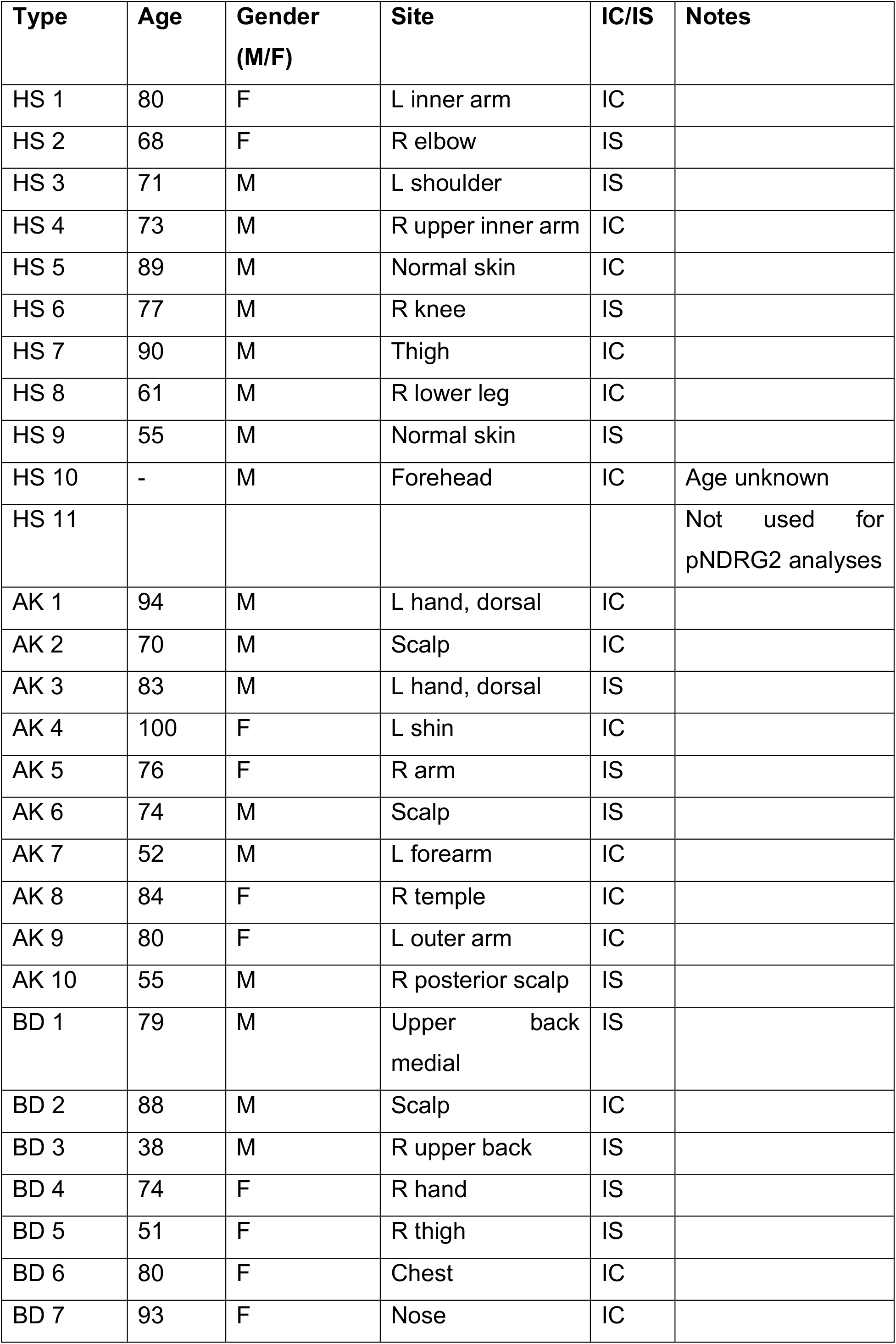
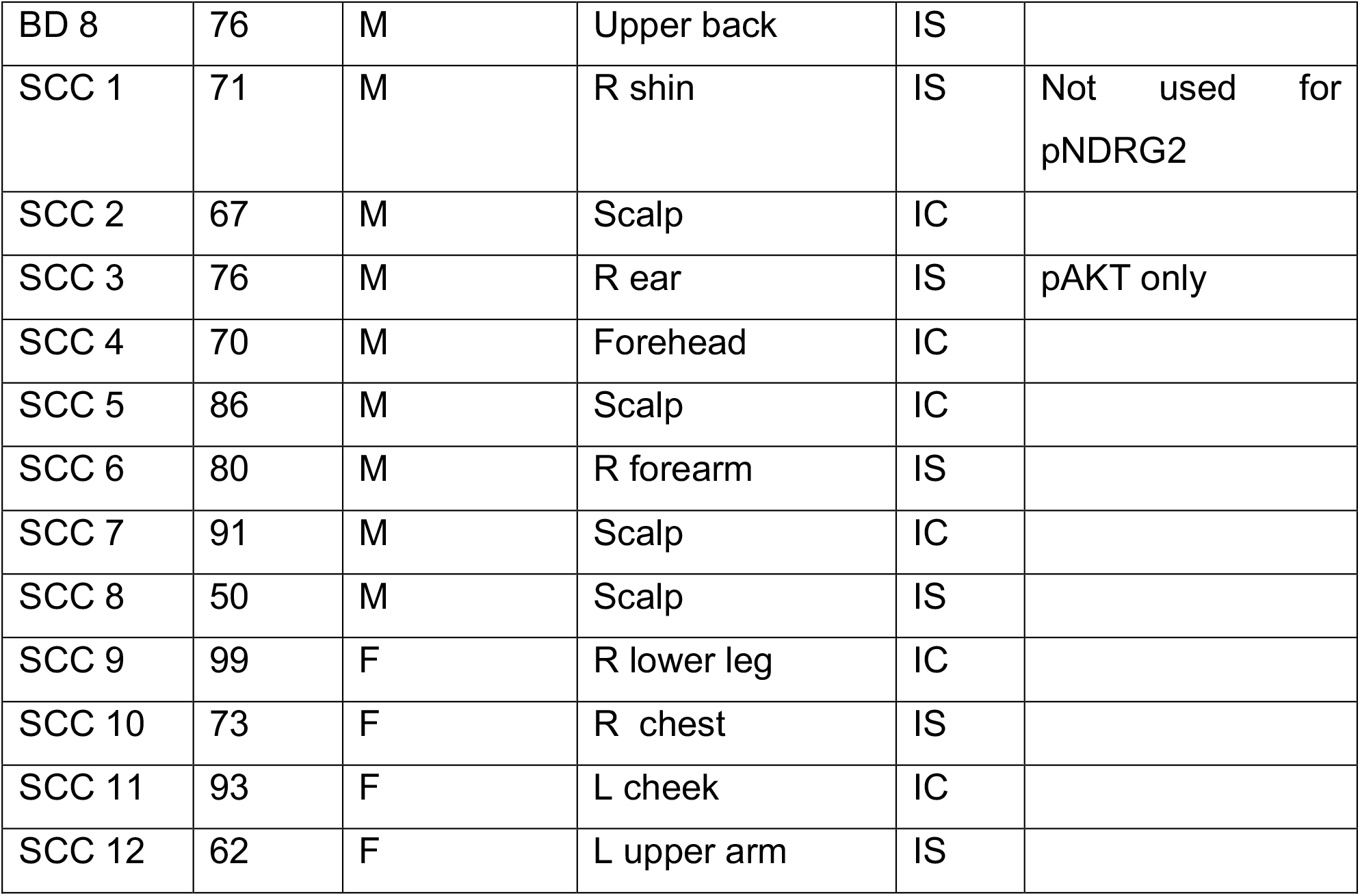
Characteristics of tumour samples used in immunofluorescence analyses. HS, normal skin; AK, actinic keratosis; BD, Bowen’s disease; SCC, Squamous Cell Carcinoma; IC, Immunocompetent; IS, Immunosuppressed.

**Supplementary Table ST2: Complete kinase set enrichment analysis data**. Heat map of significant Z-enrichment compared to pre-malignant line pM1. Red, upregulated, Blue, Downregulated. *,**,***p<0.05, p<0.005 and p<0.0005 respectively. Number of substrates and specific phosphoproteins are indicated.

**Supplementary Table ST3**: **AKT targets table**. Heat map of all putative phosphoprotein AKT targets in the complete phosphoproteomics data set. Red, upregulated, Blue, Downregulated. *,**,***p<0.05, p<0.005 and p<0.0005 respectively. Log2 Fold change compared to pM1 and corrected p-values are shown.

**Supplementary Table ST4**: **All phosphoproteins raw data**, Log2fold change and corrected p-value for comparisons of the Met1,2 and 4 lines with the pre-malignant pM1 line. Phosphorylation sites and protein descriptions are shown.

